# A co-expression network in hexaploid wheat reveals mostly balanced expression and lack of significant gene loss of homeologous meiotic genes upon polyploidization

**DOI:** 10.1101/695759

**Authors:** Abdul Kader Alabdullah, Philippa Borrill, Azahara C. Martin, Ricardo H. Ramirez-Gonzalez, Keywan Hassani-Pak, Cristobal Uauy, Peter Shaw, Graham Moore

## Abstract

Polyploidization has played an important role in plant evolution. However, upon polyploidization the process of meiosis must adapt to ensure the proper segregation of increased numbers of chromosomes to produce balanced gametes. It has been suggested that meiotic gene (MG) duplicates return to a single copy following whole genome duplication to stabilise the polyploid genome. Therefore, upon the polyploidization of wheat, a hexaploid species with three related (homeologous) genomes, the stabilization process may have involved rapid changes in content and expression of MGs on homeologous chromosomes (homeologs). To examine this hypothesis, sets of candidate MGs were identified in wheat using co-expression network analysis and orthology informed approaches. In total, 130 RNA-Seq samples from a range of tissues including wheat meiotic anthers were used to define co-expressed modules of genes. Three modules were significantly correlated with meiotic tissue samples but not with other tissue types. These modules were enriched for GO terms related to cell cycle, DNA replication and chromatin modification, and contained orthologs of known MGs. Overall 74.4 % of genes within these meiosis-related modules had three homeologous copies which was similar to other tissue-related modules. Amongst wheat MGs identified by orthology, rather than co-expression, the majority (93.7 %) were either retained in hexaploid wheat at the same number of copies (78.4 %) or increased in copy number (15.3%) compared to ancestral wheat species. Furthermore, genes within meiosis-related modules showed more balanced expression levels between homeologs than genes in non-meiosis-related modules. Taken together our results do not support extensive gene loss nor changes in homeolog expression of MGs upon wheat polyploidization. The construction of the MG co-expression network allowed identification of hub genes and provided key targets for future studies.

**Author summary:** All flowering plants have undergone a polyploidization event(s) during their evolutionary history. One of the biggest challenges faced by a newly-formed polyploid is meiosis, an essential event for sexual reproduction and fertility. This process must adapt to discriminate between multiple related chromosomes and to ensure their proper segregation to produce fertile gametes. The meiotic mechanisms responsible for the stabilisation of the extant polyploids remain poorly understood except in wheat, where there is now a better understanding of these processes. It has been proposed that meiotic adaptation in established polyploids could involve meiotic gene loss following the event of polyploidization. To test this hypothesis in hexaploid wheat, we have computationally predicted sets of hexaploid wheat meiotic genes based on expression data from different tissue types, including meiotic anther tissue, and orthology informed approaches. We have calculated homeolog expression patterns and number of gene copies for the predicted meiotic genes and compared them with proper control gene sets. Our findings did not support any significant meiotic gene loss upon wheat polyploidization. Furthermore, wheat meiotic genes showed more balanced expression levels between homeologs than non-meiotic genes.

## Introduction

Meiosis is a specialized mode of cell division which generates haploid gametes. Prior to meiosis, chromosomes are replicated. On entry into meiosis, homologous chromosomes (homologs) locate each other, and intimately align (synapse) along their length. Within this paired structure, chromosomes recombine and crossover before being accurately segregated [**1-3**]. This complex and dynamic process is essential to maintain genome stability and integrity over sexual life cycles and to generate genome variation, which is a major evolutionary driving force [**4,5**]. The genetic variation created by meiotic recombination underpins plant breeding to improve crop species [**6-8**]. Polyploidization has played an important role in the evolution and speciation of flowering plants [**9,10**], although the resultant multiplicity of related genomes poses a major challenge for the meiotic process. Segregation of the chromosomes to produce balanced gametes requires correct pairing, synapsis and recombination between only true homologs, rather than any of the other highly related chromosomes (homeologs) [**10-12**].

In the last two decades, there have been significant advances in our understanding of plant meiosis. Since the isolation of the first meiosis-specific cDNA from lily in the mid-1990s [**13,14**], more than 110 plant meiotic genes (MGs) have been identified, mainly from studies of the model diploid plants Arabidopsis and rice [**2,15,16**]. Although 25-30% of flowering plants are extant polyploids [**9**], the meiotic mechanisms responsible for their stabilisation remain poorly understood. An exception is hexaploid wheat (*Triticum aestivum* L.), where there is now better understanding of these processes [**17**]. Despite possessing multiple related genomes, durum wheat, a tetraploid (AABB) and bread wheat, a hexaploid (AABBDD) behave as diploids during meiosis. Thus, most of the meiotic studies conducted in hexaploid wheat have focused on providing better understanding of the meiotic processes required to stabilise this polyploid species [**18-22**]. An emphasis has been to characterise the role of the *Ph1* locus in the suppression of recombination between homeologs [**23-31**]. Recent studies have defined this phenotype to a *ZIP4* gene which duplicated and diverged on polyploidization [**31-33**]. This event resulted in the suppression of homeologous crossover, and promotion of homologous synapsis.

Although all flowering plants have undergone at least one event of whole genome duplication during their evolutionary history [**34**], it has been suggested that MG duplicates return to a single copy following whole genome duplication, more rapidly than the genome-wide average [**35**]. Therefore, it has been assumed that the stabilisation process upon the polyploidization of wheat also involved rapid changes in the content and expression of the genes on homeologs. This process would facilitate the correct pairing and synapsis of homeologs during meiosis. The recent development of an expression atlas for hexaploid wheat revealed that 70% of homeologous genes in syntenic triads showed balanced expression [**36**]; however, this study did not include analysis of the genes expressed during meiosis.

Here, we assessed whether the level of expression of all genes in triads was balanced between homeologs during meiosis. Analysis indicated similar balanced expression to that observed in other wheat tissues. However, it could be argued that only meiotic specific genes might show differential expression between homeologs. Sets of candidate MGs were identified using co-expression network analysis and orthology informed approaches, allowing us to evaluate the effect of polyploidization on wheat MG copy number and expression. The combination of co-expression network analysis, in conjunction with orthologue information, will now contribute to the discovery of new MGs and greatly empower reverse genetics approaches to validate the function of candidate genes in wheat.

## Results and Discussion

An initial assessment of the homeolog expression pattern in triads during meiosis in hexaploid wheat was undertaken. Relative expression abundance of 19,801 triads (59,403 genes) was calculated for 8 tissues, including meiotic anther tissue, according to published criteria [**36**]. This analysis revealed that the percentage of balanced triads was slightly higher in meiotic anther tissue (77.3%) than in other type of tissues (ranging from 67.3% in floral organs and 76.6% in leaves) (**S1 Fig**). The copy number of genes expressed during meiosis was also investigated. This involved the definition of 19801 triads (59403 genes), 7565 duplets (15130 genes), 15109 monads (single-copy genes) and 18250 genes from the “Others” group with various copy numbers, based on the *Ensembl*Plants database for the HC genes of hexaploid wheat [**37**] (IWGSC v1.1 gene annotation; **S1 Table**). Comparison of copy number of genes expressed in the 8 different tissues showed that 70.9% of the genes expressed during meiosis belonged to triads. This percentage ranged between 66.5% and 72.5% for the genes expressed in floral organs and stem tissues, respectively (**S2 Fig**). These results were consistent with a previous study reporting significant balanced expression between homeologous genes in tissues other than meiotic anthers [**36**]. However, the observations were not in agreement with the hypothesis that stabilisation of polyploidization involved significant changes in gene content and expression between homeologs [**35,38,39**]. Considering that not all genes expressed in meiotic anther tissue are directly involved in the meiosis process, it is possible that meiotic specific genes exhibit a different pattern. Therefore, a co-expression gene network was developed to compare the expression pattern of homeologous genes in meiosis-related modules, which potentially represent meiosis specific genes, and other tissue-related modules.

### Weighted co-expression network construction

Network-based approaches have been proved useful in systems biology, to mine gene function from high-throughput gene expression data. Gene co-expression analysis has become a powerful tool to build transcriptional networks of genes involved in common biological events in plants [**40-45**]. The use of co-expression networks has uncovered candidate genes to regulate biological processes in many plants including wheat [**37**], rice [**46,47**] and *Arabidopsis thaliana* [**48,49**]. The recently published high-quality genome reference sequence [**37**] and a developmental gene expression atlas [**50,36**], together with the gene expression data collected from meiotic samples, were used to build a co-expression gene network. One hundred and thirty samples from different tissue types were included in this co-expression analysis (**S2 Table**; **Fig 1A**). A set of 60,379 genes out of the total 107,892 HC genes was considered expressed during meiosis (transcript per million (TPM) > 0.5 in at least one meiosis sample) and used to run the co-expression analysis. Using the “WGCNA” package in R [**51,52**], genes with similar expression patterns were grouped into modules via the average linkage hierarchical clustering of normalized count expression values (**Fig 1E**). The power of β = 7 (scale free topology R^2^ = 0.91) was selected as the soft threshold power to emphasize strong correlations between genes and penalize weak correlations to ensure a scale-free network (**Fig 1B-C**). Based on this analysis, 50,387 out of 60,379 genes (83.5% of expressed genes) could be assigned to 66 modules. Module size ranged from 52 to 7541 genes (mean 763 genes; median 429 genes) (**Fig 1D**). Detailed information about gene number and gene IDs in each module is shown in **S3 Table**.

**Fig 1.**
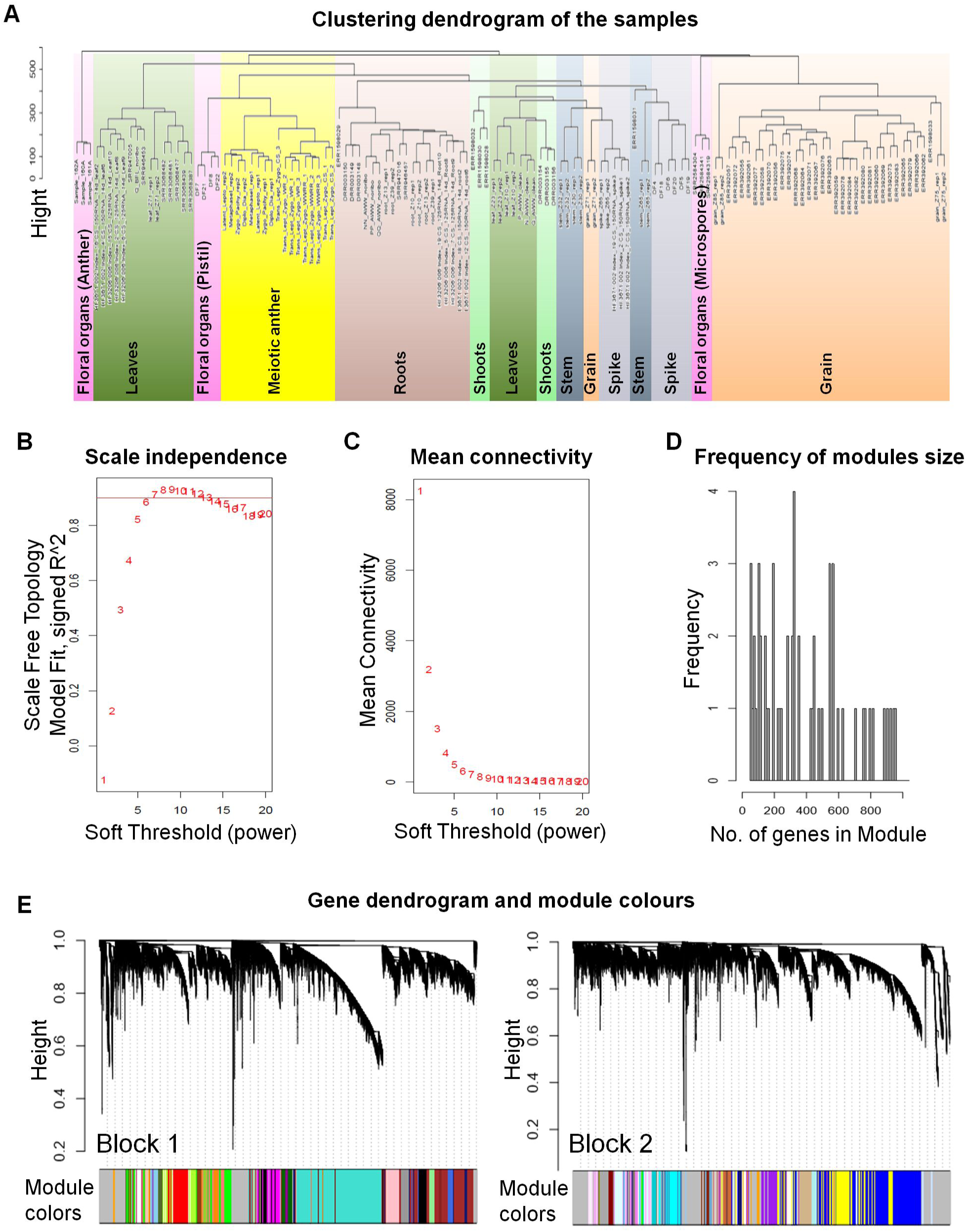
The weighted gene co-expression networks analysis (WGCNA). **(A)** Clustering dendrogram of 130 samples from different types of tissues. The sample clustering was based on the expression data of the genes expressed in at least one meiosis sample. **(B)** Determination of soft-thresholding power (β). The soft power threshold used in constructing the weighted gene co-expression networks was chosen as the first power to exceed a scale-free topology fit index of 0.9 (then β = 7). **(C)** Analysis of the mean connectivity for various soft-thresholding powers. **(D)** Number of genes in the modules with their frequency. **(E)** Blockwise dendrogram of the analysed genes (60,379 genes) clustered based on a dissimilarity measure of topological overlap matrices (TOM).

The expression patterns of all genes within a single module were summarized into a Module Eigengene (ME; representative gene of the module) to minimize data size for subsequent analyses. Expression patterns of modules are shown as a heatmap by plotting ME values in relation to tissue samples (**Fig 2**).

**Fig 2.**
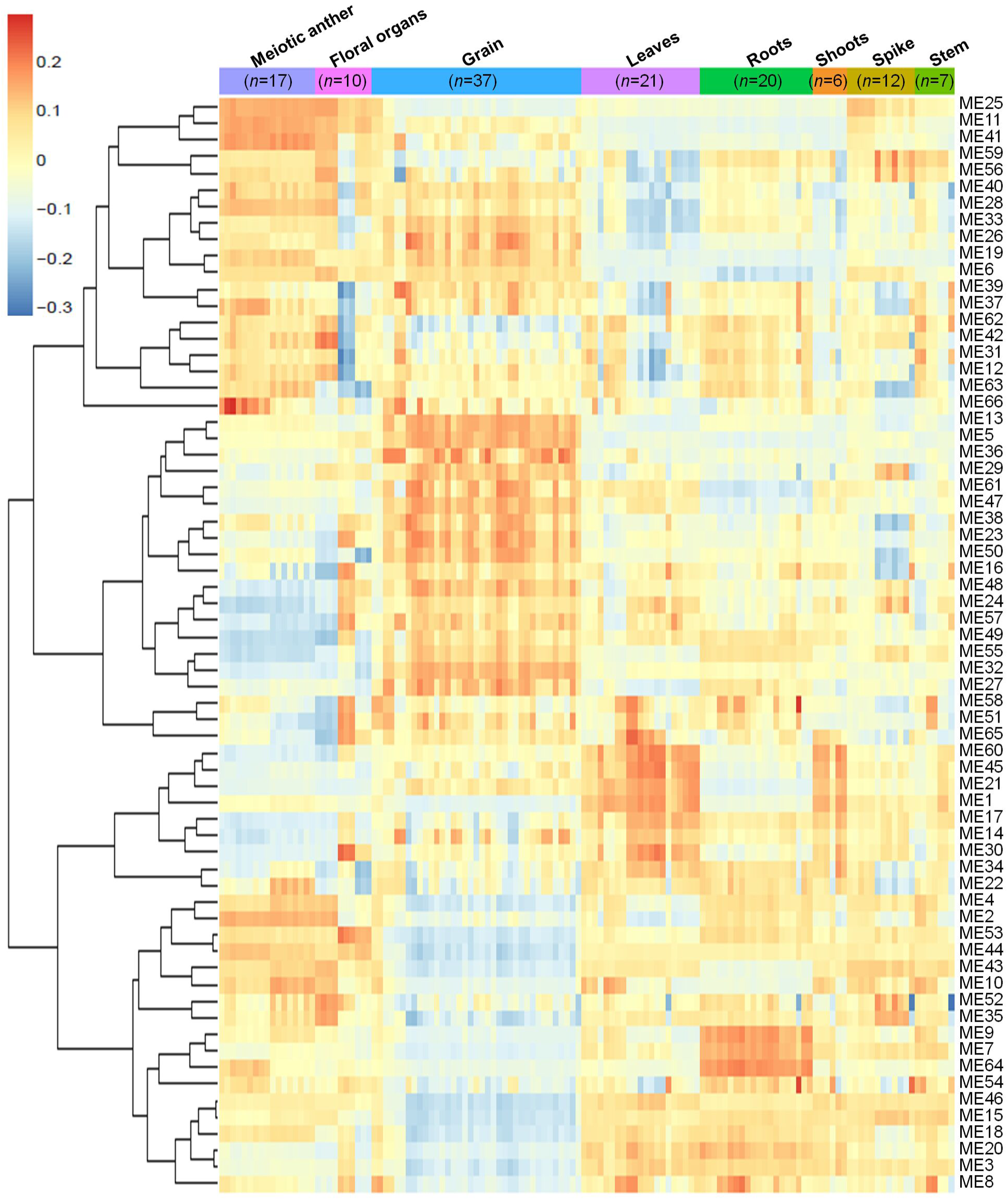
Heatmap plotting of MEs values in relationship to tissue samples. *n* indicates number of samples.

### Identification of meiosis-related modules

A correlation analysis was conducted using the 66 MEs and the 8 different tissue types. A module was considered as meiosis-related when there was a strong correlation (*r*) with the 17 meiosis samples, and a weak or negative correlation with other tissue types. Accordingly, three meiosis-related modules were identified: module 2 (containing 4,940 genes); module 28 (544 genes); and module 41 (313 genes). Module 41 showed the strongest correlation with meiotic tissue (*r* = 0.73, FDR =2.7 × 10^−20^), compared to module 2 (*r* = 0.61, FDR =9.2 × 10^−13^) and module 28 (*r* = 0.52, FDR =2.1 × 10^−8^). (**Fig 3**).

**Fig 3.**
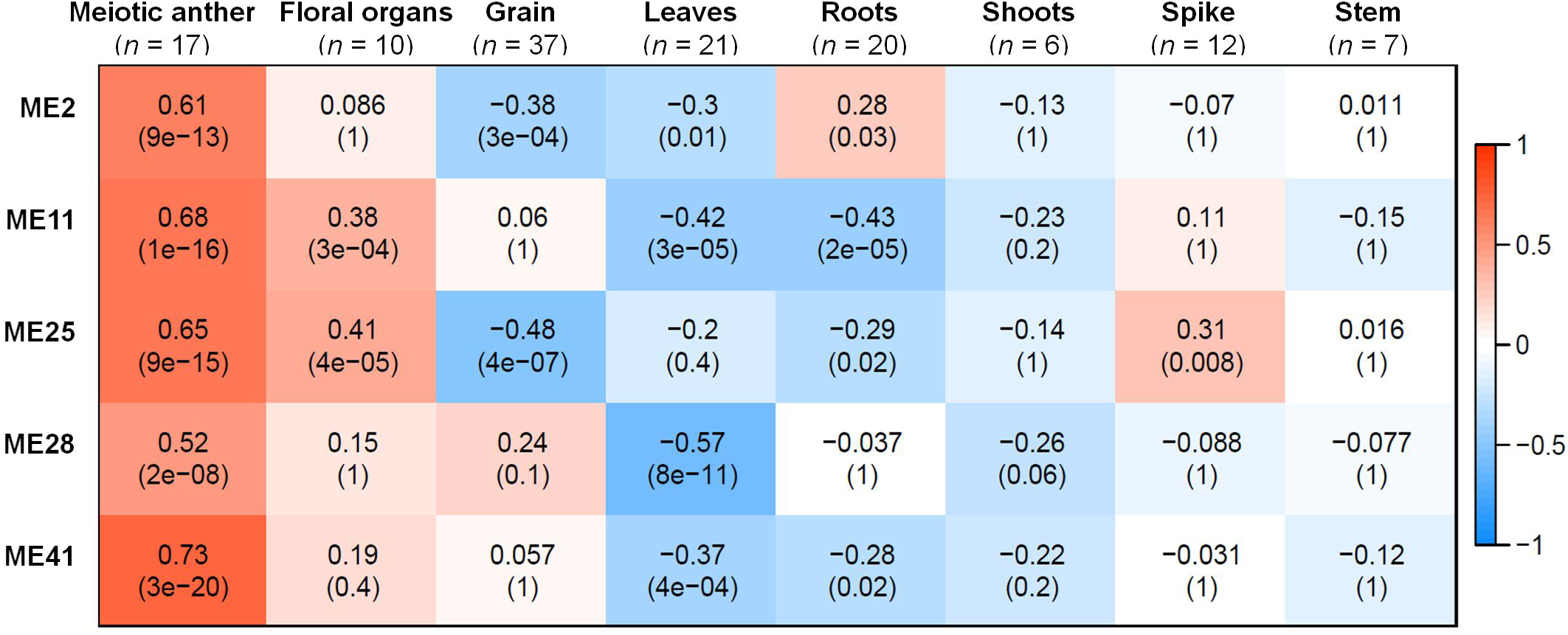
Co-expression network modules in relationship to tissues samples. Each row corresponds to a module; each column corresponds to a tissue type; Each cell contains the correlation value and, in parentheses, its corresponding FDR adjusted *P* value. *n* indicates number of samples. Only modules that have correlation value > 0.5 with meiotic anther tissue are shown.

Two other modules (modules 11 and 25) also showed significantly positive correlation with meiotic samples (*r* = 0.68 and 0.65, respectively), however they were not considered meiosis-related because they also correlated with samples from floral organs (at stages other than meiosis) and spike tissues, as shown in **Fig 3**. Therefore, our analysis focused on the three modules (2, 28 and 41), exhibiting a strong correlation with meiotic tissues and not with other floral organs, while modules 11 and 25 were considered as non-meiosis specific modules (referred to in this paper as non-meiotic modules). Other tissue-related modules (the top three correlated modules) were also identified to be used as controls for the meiosis-related modules in the subsequent analysis. These modules were: grain-related modules 5, 13 and 32 (*r* = 0.89, 0.89 and 0.85, respectively); leaves-related modules 1, 45 and 60 (*r* = 0.72, 0.68 and 0.71, respectively); and roots-related modules 7, 9 and 64 (*r* = 0.70, 0.76 and 0.85, respectively) **(S3 Fig)**.

### Biological significance of expression similarity in modules

Several approaches were undertaken to validate the meiosis-related modules. The three modules (2, 28 and 41), strongly correlated with meiotic tissue expression, were found to be significantly enriched with the gene ontology (GO) slim terms “cell cycle”, “DNA metabolic process”, “nucleobase-containing compound metabolic process” and “nucleus” (**Fig 4**). Among the top five enriched GO slim terms in each of the 66 modules, the term “cell cycle” was significant only in the three meiosis-related modules, suggesting this was not a general property of all modules, and was instead specific to the meiosis modules. Module 2 in particular was significantly enriched with GO terms related to many biological processes occurring during meiosis such as “DNA replication”, “histone methylation”, “cytokinesis”, “nucleosome assembly” and “chromatin silencing” (**Table 1; S4 Table)**. The term “double-strand break repair via homologous recombination”, an important process during meiosis, was the primary enriched Biological Processes GO term in module 41 (FDR < 0.05). The biological processes mediated by genes in module 28 included “protein deneddylation”, “positive regulation of G2/M transition of mitotic cell cycle”, “COP9 signalosome” and other terms related to protein deneddylation and cell cycle control (**Table 1**). GO terms of meiosis-related modules were compared with those of modules highly correlated with other tissues. The GO terms “chloroplast”, “plastids”, “thylakoid”, “photosystem” were significantly enriched in module 1, the most highly correlated module with leaves. The terms related to protein ubiquitination and protein binding were enriched in module 5 (the most highly correlated module with grain), while the terms “lignin biosynthetic process”, “phenylpropanoid metabolic process” and “response to wounding” were enriched in module 7, the largest module correlated with roots (**S4 Fig**). This indicated that our co-expression module-tissue correlation was meaningful both from the biological and physiological point of view. Detailed information of the enriched GO and GO slim terms in all modules is listed in the supplementary table **S4 Table**. In summary, GO analysis confirmed that the three modules (2, 28 and 41) were enriched for genes associated with meiotic processes.

**Fig 4.**
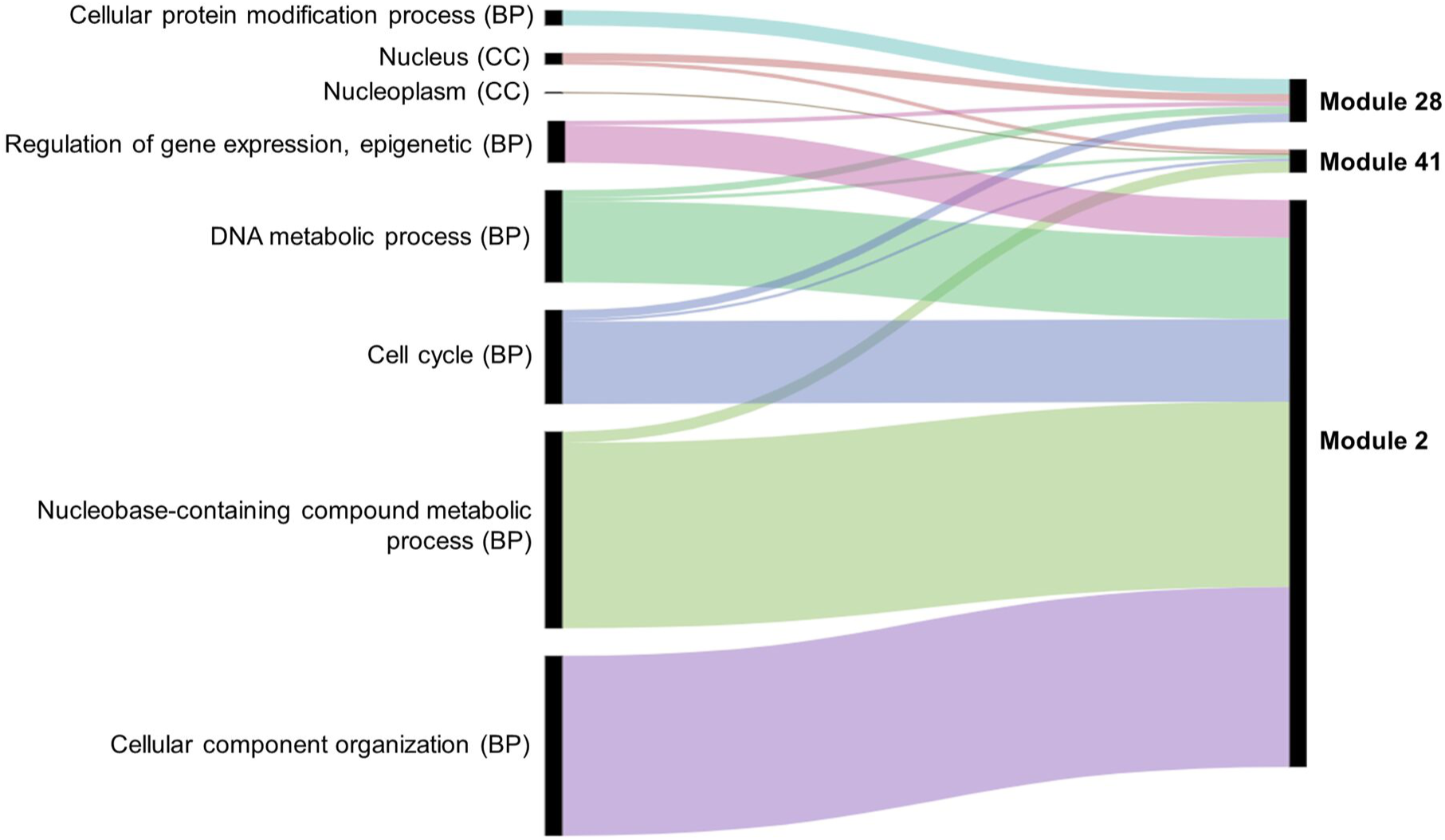
Enriched GO slim terms in the meiosis-related modules. Top 5 enriched GO terms in each module are shown. BP indicates Biological Processes and CC Cellular Component. No Molecular Function (MF) GO terms appear among the top 5 GO slim terms. Black bars indicate the number of genes in the GO term.

**Table 1.**
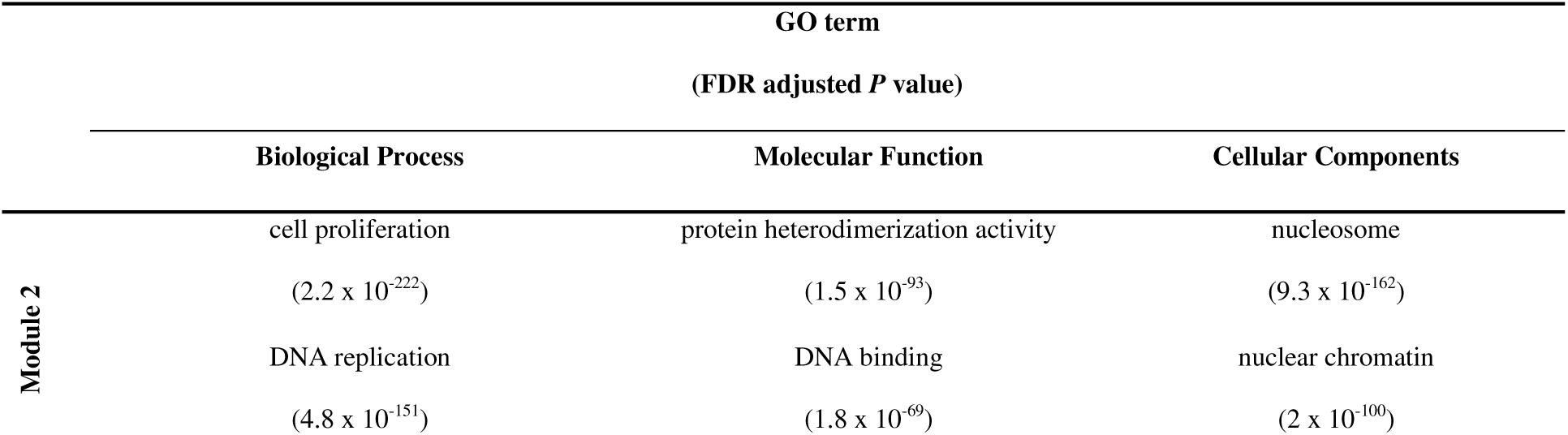
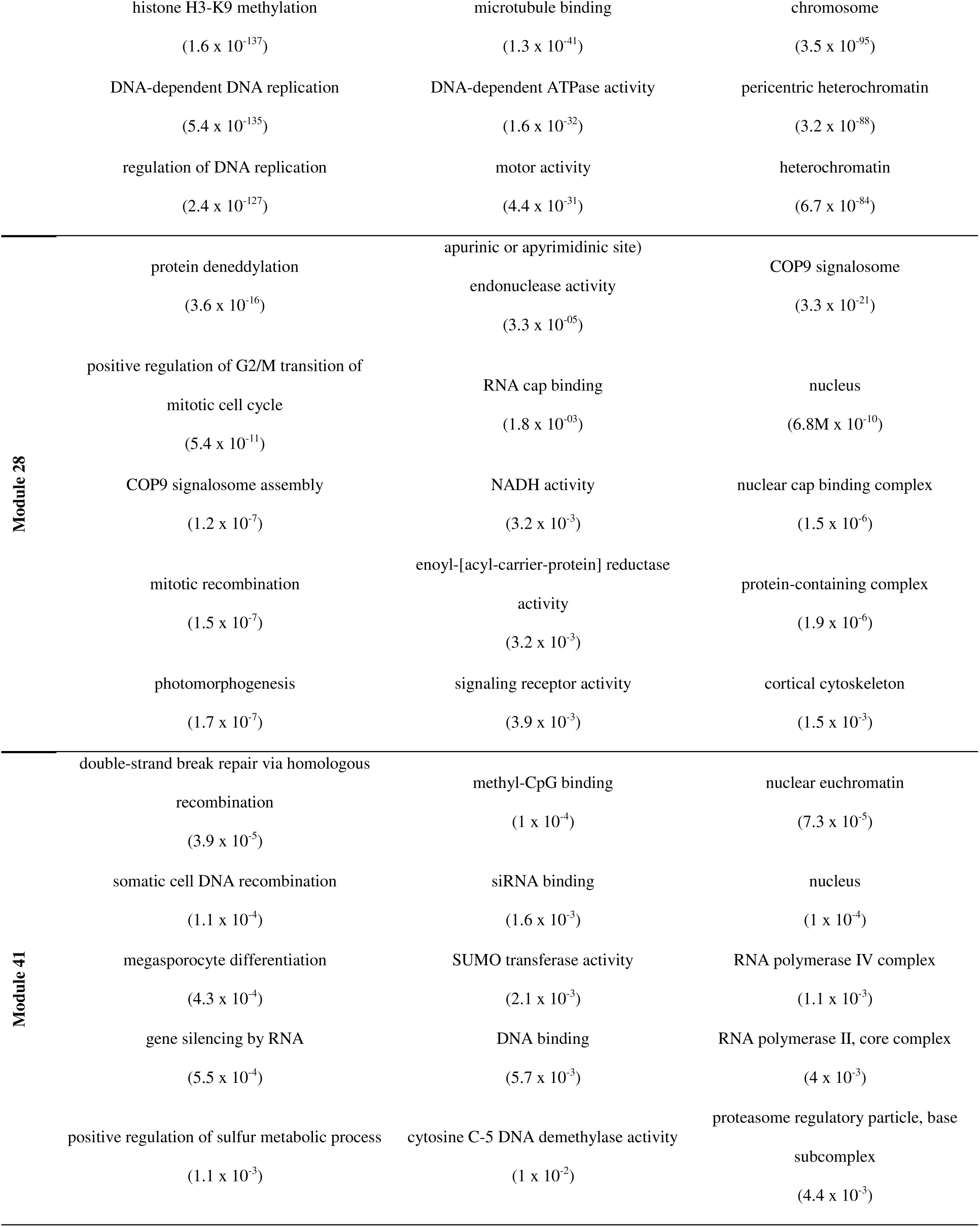
Top 5 enriched GO terms in the meiosis-related modules for each ontology group.

### Enrichment of meiosis-related modules for wheat orthologs of known MGs

An assessment was undertaken to confirm that the meiosis-related modules were enriched for wheat orthologs of known MGs. Although the first wheat meiotic cDNA clones were isolated concurrently with the early discoveries of MGs in other plants [**53**], the identification of MGs in wheat has been hampered by the large wheat genome size, its polyploid nature and the absence of a complete genome sequence. Thus, in comparison to model plants (Arabidopsis and rice), few MGs have been functionally characterized in wheat. Characterised wheat MGs include *TaASY1* [**54,55**], *TaMSH7* [**56**], *TaRAD51* [**57,58**], *TaDMC1* [**57,58**], *TaPSH1* [**59**], *TaZIP4* [**31-33**] and RecQ-7 [**60**]. Given that, the assessment of whether the three modules contain known MGs was undertaken using orthology informed approaches. A set of 1063 candidate MGs in wheat was identified and categorized based on the method used to identify the genes: the “Orthologs” group contained 407 genes (**S5 Table; S6 Table)** that correspond to wheat orthologs of 103 functionally characterized MGs in model plant species, and the “Meiotic GO” group contained 927 wheat genes annotated with one or more meiotic GO terms (**S7 Table**). There were 271 genes overlapping between the two groups (**Fig 5A**), that were considered in the “Orthologs” group when undertaking gene enrichment analysis. The presence of each gene in the different modules was determined. A set of 848 genes was assigned to modules in the co-expression network (**Fig 5B**), including 340 genes in meiosis-related modules. Genes from both groups were significantly over-represented (*P* < 0.05) in four modules, including two meiosis-related modules (2 and 28). Module 2 in particular, was the most enriched for these genes, possessing more than one third of the total candidate MGs assigned to modules. Module 2 had 142 wheat orthologs of MGs and 155 genes with meiotic GO terms, compared to the expected number (based on module size) of 27 and 42, respectively. Module 41 (the third meiosis-related module) was enriched only with genes from the “Orthologs” group, having 15 orthologs of known MGs, whereas the expected number was 2 (**Fig 5C**). Consistent with this, genes from the “Orthologs” and/or “Meiotic GO” groups, were significantly under-represented in modules strongly correlated with other types of tissue (modules 1, 7 and 9), and in modules with negative or no correlation with meiotic tissue (modules 3, 8, 14, and 17) (**S8 Table**).

**Fig 5.**
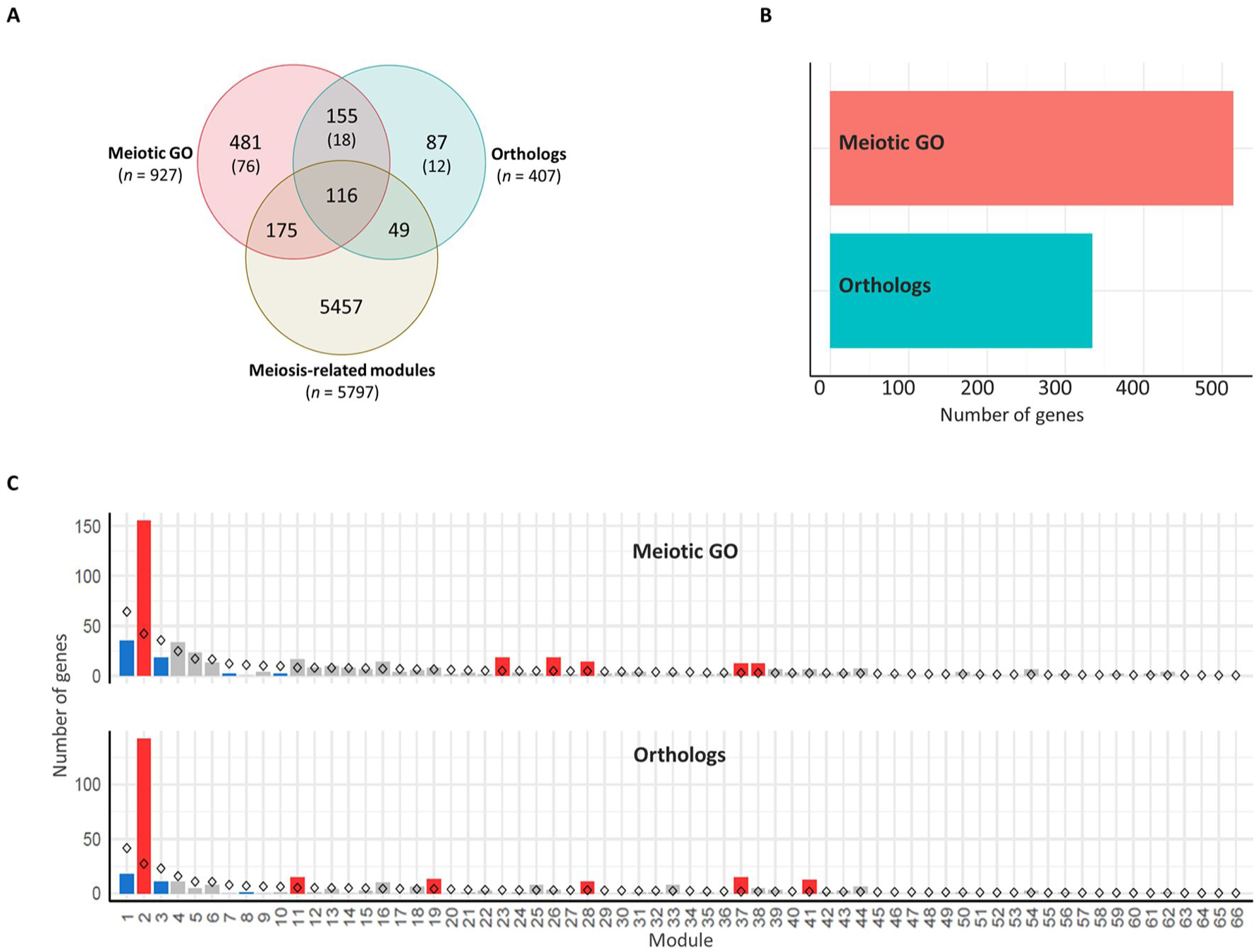
Enrichment of meiosis-related genes in the co-expression network modules. (**A**) Venn diagram of overall genes in the meiosis-related modules (modules 2, 28 and 41), wheat orthologs of MGs in model plant species and genes with meiotic GO terms. *n* indicates number of genes in each group. Numbers in brackets refer to number of genes not included in the WGCNA analysis because they are not expressed in meiotic anther tissue. (**B**) Total number of genes assigned to modules from orthologs of MGs and genes with meiotic GO terms. (**C**) Gene enrichment in modules. Statistical significance of gene enrichment in a module is colour coded (Red indicates over-represented, blue under-represented and grey not significant; *P* < 0.05). Rhombus shape indicates the expected number of genes in module.

In this study three co-expression gene modules were identified that are strongly correlated to meiotic anther tissue and highly enriched with GO terms related to many processes occurring during meiosis, orthologs of known MGs and genes having meiosis-specific GO terms. Although 67 (65%), out of the 103 wheat orthologs, had at least one gene copy assigned to one of the three meiosis-related genes, there were 36 orthologs whose gene homeologs were assigned to other modules (**S9 Table**). Some of those genes have essential meiotic functions, like: *ASY1*, which encodes a protein essential for homologous chromosome synapsis [**54,61,62**]; and *DMC1*, a gene encoding a recombination protein that acts only in meiosis [**58,63**]. Others are known to have both meiotic and mitotic functions, like: *BRCA2*, a DNA repair gene required for double strand breaks repair by homologous recombination [**64**] and *SMC1* and *SMC3*, chromosome cohesion genes [**65**], thus they are expressed in both reproductive and non-reproductive tissues. Assessment of these 36 orthologs showed that the expression patterns of their gene copies did not allow them to be clustered in any of the meiosis-related modules (or allocated to module 0 that is composed of genes not forming part of a co-expressed module), either because they were expressed in most samples from all types of tissues, or because they were expressed in a few samples of a specific tissue type (like meiotic anther tissue). The expression values (TPMs) of all gene copies of those 36 orthologs are summarized in **S10 Table**. The number of meiotic anther samples (17) used in the present study, might not be enough to identify all MGs being expressed in a specific meiotic stage. Such genes might be identified by WGCNA analysis when a larger number of meiotic samples becomes available. However, the analysis confirmed that the meiosis-related modules were indeed enriched for orthologs of known MGs, and for GO terms associated with processes involved in meiosis.

### Copy number of MGs

It has previously been suggested that MG duplicates return to a single copy following whole genome duplication more rapidly than the genome-wide average in angiosperms [**35**]. The analysis of 19 meiotic recombination genes in hexaploid wheat and oilseed rape showed no evidence of gene loss after polyploidization. However, a recent study in tetraploid oilseed rape showed that reducing the copy number of *MSH4*, a key meiotic recombination gene involved in the ZMM pathway, prevents meiotic crossovers between non-homologous chromosomes [**66**]. This led to the suggestion that meiotic adaptation in polyploids could involve ‘fine-tuning’ the progression or the effectiveness of meiotic recombination, which could be achieved through the loss of MG duplicates in the newly-formed polyploids [**35,66**]. This hypothesis was evaluated in hexaploid wheat. The gene copy number was assessed for the genes in the three meiosis-related modules and compared with genes in all modules and in other tissue-related modules. Analysis showed that the percentage of genes belonging to triads was 74.4% in the meiosis-related modules, which was similar to this percentage in other tissue-related modules (72.5%, 74.0% and 76.1% in leaves-, grain- and roots-related modules, respectively); however, it was significantly higher than those of the non-meiotic modules (57.4%). The highest percentage of genes with three homeologs (83.3%) and lowest percentage of genes with single copy (2.7%) were observed in the group of genes identified as MG orthologs and/or possessing a meiotic GO term (**Fig 6A**).

**Fig 6.**
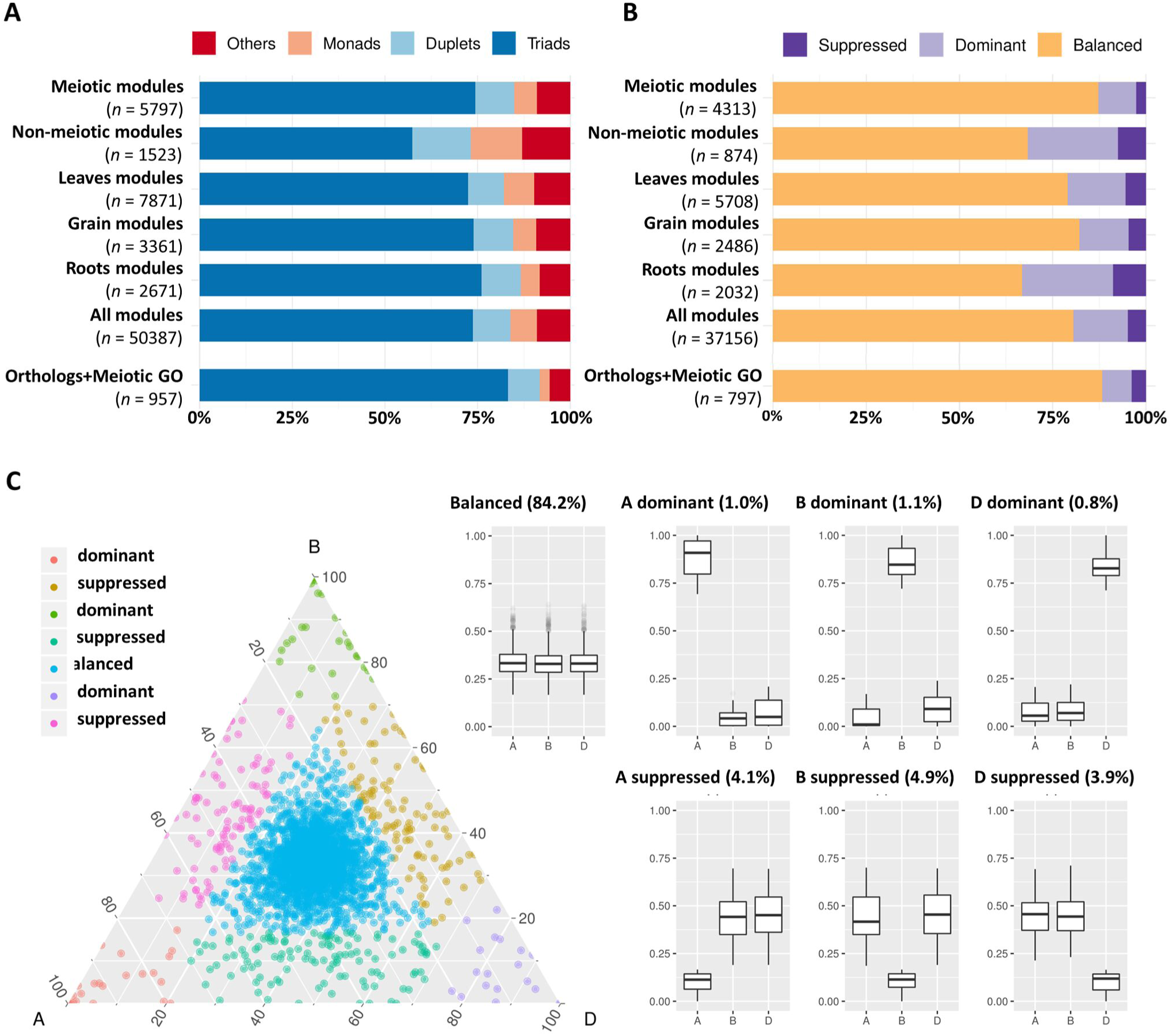
Copy number and homeolog expression pattern for genes from meiosis-related and other tissue-related modules. (**A**) Proportion of genes in each copy number category (triads, duplets, monads and others) for different sets of expressed genes during meiosis including: “Meiotic modules” refers to the three meiosis-related modules 2, 28 and 41; “Non-meiotic modules” refers to the modules 11 and 25 that showed high correlation with meiotic anther but were not considered meiosis-related because they were also correlated with spike and floral organs tissues; the top 3 correlated modules with each of leaves (modules 1, 45 and 60), grain (modules 5, 13 and 32) and roots (modules 7, 9 and tissues; “All modules” contains all genes assigned to modules in the co-expression network and “Orthologs & Meiotic GO” refers to the set of genes that are orthologs of known MGs in other plant species and/or have meiotic GO terms. *n* number of genes in each set. (**B**) Proportion of genes from each homeolog expression pattern category (balanced, dominant and suppressed) calculated for triads in the previously mentioned sets of genes. *n* number of genes in each set. (**C**) Ternary plot showing relative expression abundance in meiotic anther tissue of 2,366 triads to which the genes of meiosis-related modules (2, 28 and 41) belong. Each circle represents a gene triad with an A, B, and D coordinate consisting of the relative contribution of each homeolog to the overall triad expression. Triads in vertices correspond to single-subgenome-dominant categories, whereas triads close to edges and between vertices correspond to suppressed categories. Box plots indicate the relative contribution of each subgenome based on triad assignment to the seven categories (Balanced, A dominant, B dominant, D dominant, A suppressed, B suppressed, D suppressed). Percentages between brackets indicate the percentage of triad number in each category to the total number of triads.

The high percentage of meiosis-related genes present as triads provides evidence that polyploid wheat did not experience significant gene loss (gene erosion) after polyploidization. However, this assumes that these genes were originally present as single copy genes in each of the A-, B- and D-genome progenitor species which gave rise to polyploid wheat. Therefore, the copy number of the 103 wheat MG orthologs in wheat progenitor species was investigated. All possible orthologs (high and low confidence predicted orthologs) were retrieved from *Ensembl* Plants Genes 43 database for *Triticum urartu* (ASM34745v1; [**67**]), the diploid progenitor of the wheat A-genome, the D-genome ancestor *Aegilops tauschii* (Aet_v4.0; [**68**]), the diploid progenitor of the wheat D-genome and *Triticum dicoccoides* (WEWSeq_v.1.0; [**69**]), the tetraploid progenitor of the hexaploid wheat (genome AABB). There was no change in copy number of 78.4% of genes, while 6.3% and 15.3% of genes had a lower and greater number of copies, respectively (**Table 2** and **S11 Table**). Regardless of genome of origin, the percentage of MGs with more copies was always greater than the percentage of genes with fewer copies. Comparing the A-genome MGs copy number in hexaploid wheat with the relevant orthologs copy number in the corresponding A-genome ancestor, 86 genes (84.5%) had the same gene copy number in *T. dicoccoides* as in hexaploid wheat, while only 64 genes (63.1%) had the same gene copy number in *T. urartu*. This is consistent with the evolutionary history of hexaploid wheat, with *T. dicoccoides* being a more recent wheat progenitor (∼10,000 years) than *T. urartu* (> 5 million years) [**70**].

**Table 2.**
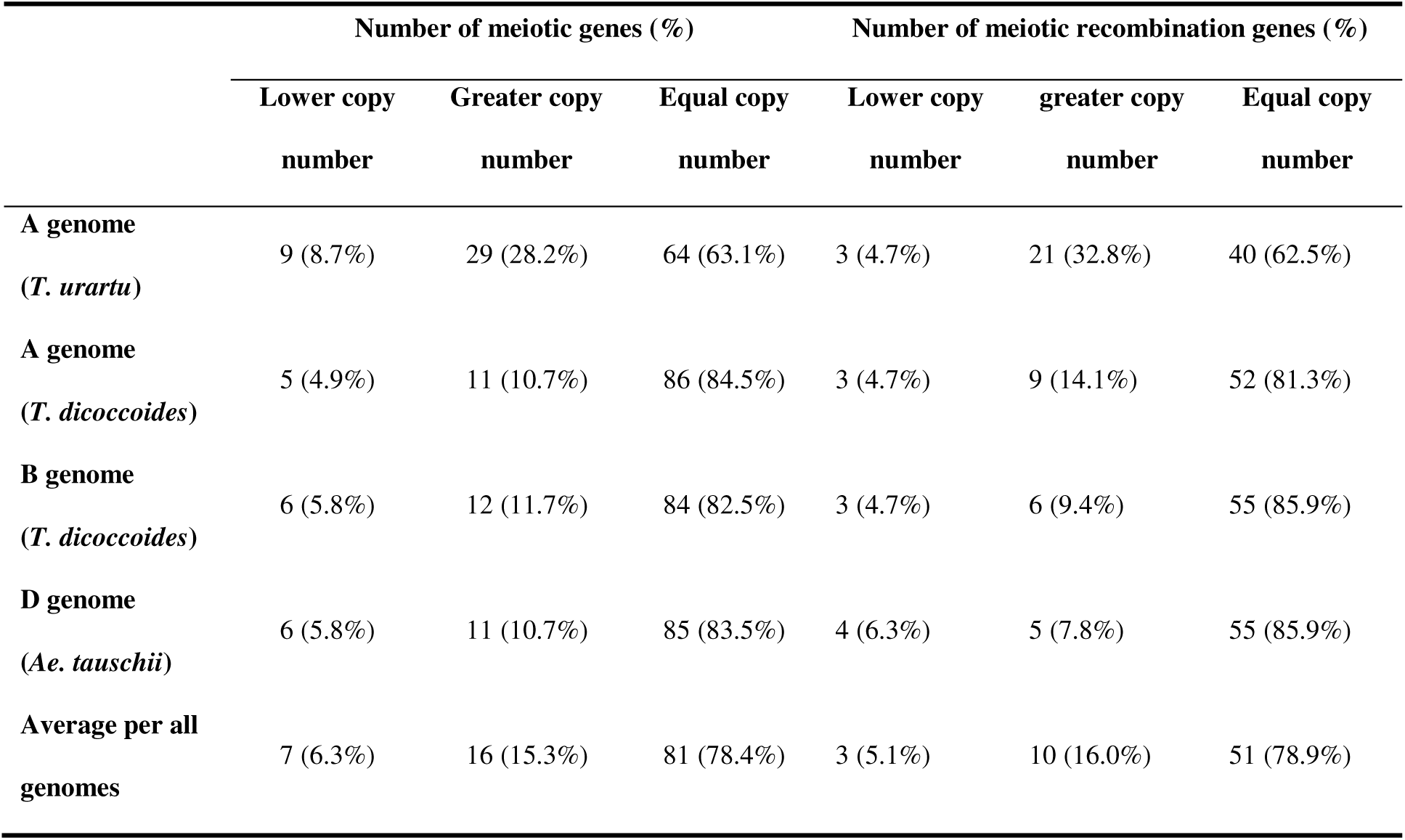
Changes in copy number of wheat MGs in comparison with their orthologs in wheat progenitors. Comparison was done for two sets of genes: wheat orthologs of MGs (*n* = 103) and wheat meiotic recombination genes (*n* = 64).

|  | Number of meiotic genes (%) |  |  | Number of meiotic recombination genes (%) |  |  |
| --- | --- | --- | --- | --- | --- | --- |
|  | Lower copy | Greater copy | Equal copy | Lower copy | greater copy | Equal copy |
|  | number | number | number | number | number | number |
| **A genome**<br>(*T. urartu*) | 9 (8.7%) | 29 (28.2%) | 64 (63.1%) | 3 (4.7%) | 21 (32.8%) | 40 (62.5%) |
| **A genome**<br>(*T. dicoccoides*) | 5 (4.9%) | 11 (10.7%) | 86 (84.5%) | 3 (4.7%) | 9 (14.1%) | 52 (81.3%) |
| **B genome**<br>(*T. dicoccoides*) | 6 (5.8%) | 12 (11.7%) | 84 (82.5%) | 3 (4.7%) | 6 (9.4%) | 55 (85.9%) |
| **D genome**<br>(*Ae. tauschii*) | 6 (5.8%) | 11 (10.7%) | 85 (83.5%) | 4 (6.3%) | 5 (7.8%) | 55 (85.9%) |
| **Average per all genomes** | 7 (6.3%) | 16 (15.3%) | 81 (78.4%) | 3 (5.1%) | 10 (16.0%) | 51 (78.9%) |

An analysis on a subset of wheat genes, which were expected to be involved in meiotic recombination based on the function of their orthologs in model plants (64 genes; **S11 Table**), was conducted. Again, results showed that the majority (94.9%) of those genes had greater or no change in number of copies (**Table 2**). Given it has been suggested that the reduction in the copy number of ZMM pathway genes could stabilize meiosis in Brassica [**66**], the copy number of the wheat orthologs of seven ZMM genes was evaluated. Five of the seven ZMM genes (*MER3*, *MSH5*, *ZIP4*, *PTD* and *SHOC1*) had equal or greater number of copies. However, *TaMSH4* gained one A-genome copy (comparing with *T. urartu*) and lost one D-genome copy (compared with *Ae. tauschii*), while *TaHEI10* lost A-genome copy and gained a D-genome copy (**S11 Table)**. In conclusion, our findings did not support any significant gene loss upon the polyploidization of hexaploid wheat, as suggested for other polyploids [**35,66,71**].

### Homeolog expression patterns in triads of MGs

Initial analysis revealed that most genes expressed during meiosis showed balanced expression between homeologs (**S1 Fig**). The analysis was repeated using gene expression within the validated meiosis modules. Genes from all modules were assigned to three categories (balanced, dominant and suppressed) (see Materials and Methods; **S12 Table**). Homeolog expression patterns in triads showed that meiosis-related modules 2, 28 and 41 had the highest percentage (87.3%) of genes with balanced expression (belong to balanced triads), compared to the top three tissue-related modules for grain, leaves and roots (**Fig 6B**). Surprisingly, the group of candidate MGs selected for being orthologs of known MGs in other plant species and/or having meiotic GO terms had a higher percentage of genes from balanced triads (88.3%); whereas the modules not considered meiosis-specific (having high correlation with meiotic anther tissue and with spike and floral organs) contained only 68.3% genes with balanced expression (**Fig 6B**). The majority (84.19%) of triads with genes in meiosis-related modules (2,366 triads) showed balanced expression in meiotic anther tissue (**Fig 6C**).

In wheat, meiotic recombination and gene evolution rates are strongly affected by chromosome position, with relatively low recombination rates in the interstitial and proximal regions (genomic compartments R2a, C and R2b) but notably higher rates toward the distal ends of the chromosomes (genomic compartments R1 and R3) [**72,73**]. The lack of significant changes in gene content and more balanced expression between homeologs, suggested that these genes might be more prevalent in the proximal genomic compartments [**36,37**]. The distribution of MGs was therefore assessed across the genomic compartments compared with the distribution of all HC genes across chromosomes (**S13 Table**). Analysis showed that genes from the meiotic modules (modules 2, 28 and 41), were significantly over-represented in the genomic compartments R2a, C and R2b (*P* = 2.4 × 10^−5^, 3.1 × 10^−6^ and 1 × 10^−5^, respectively), while they were under-represented in the R1 and R3 genomic compartments (*P* = 1.7 × 10^−8^ and 1.7 × 10^−10^, respectively) (**Fig 7A**). Enrichment in the R2a genomic compartment region was not observed for genes from any of the other top three tissue-related modules (**Fig 7B**), since 21.7% of genes from the meiosis-related modules were assigned to R2a, while this percentage ranged between 18.2% and 19.5% in other tissue-related modules (**Table 3**). Interestingly, the set of the genes identified through orthology approaches had also similar high percentage (21%) of genes assigned to R2a genomic compartment (**Table 3**).

**Fig 7.**
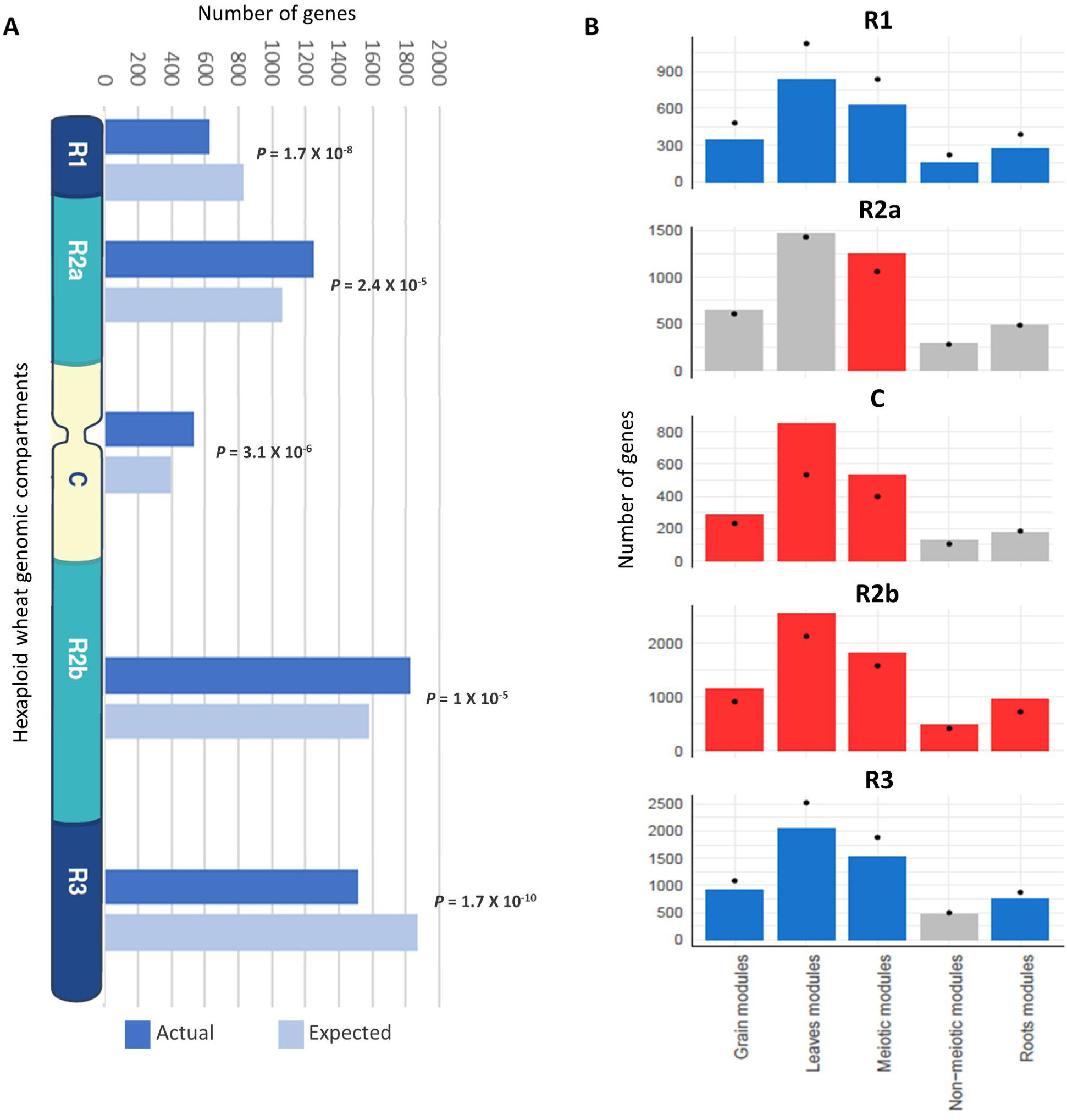
Enrichment of genes from different tissue-related modules in the wheat genomic compartments. **(A)** Number of genes (actual and expected) from the three meiosis-related modules in each genomic compartment. **(B)** Comparison of number of genes from different tissue-related modules in genomic compartments. Statistical significance of gene enrichment in modules is colour coded (Red indicates enriched, blue depleted and grey not significant; *P* < 0.05). Black dots indicate the expected number of genes in groups.

**Table 3.** Number of genes from different groups in the wheat genomic compartments.

| Modules | R1 |  | R2a |  | C |  | R2b |  | R3 |  |
| --- | --- | --- | --- | --- | --- | --- | --- | --- | --- | --- |
|  | No. | % | No. | % | No. | % | No. | % | No. | % |
| Meiotic modules | 625 | 10.9% | 1246 | 21.7% | 531 | 9.3% | 1823 | 31.8% | 1515 | 26.4% |
| Non-meiotic modules | 158 | 10.5% | 292 | 19.5% | 123 | 8.2% | 469 | 31.3% | 456 | 30.4% |
| Grain modules | 338 | 10.2% | 649 | 19.5% | 287 | 8.6% | 1144 | 34.4% | 903 | 27.2% |
| Leaves modules | 838 | 10.8% | 1466 | 18.9% | 853 | 11.0% | 2557 | 33.0% | 2037 | 26.3% |
| Roots modules | 270 | 10.3% | 479 | 18.2% | 177 | 6.7% | 951 | 36.1% | 754 | 28.7% |
| All modules | 5147 | 10.3% | 9880 | 19.9% | 4507 | 9.1% | 16491 | 33.1% | 13738 | 27.6% |
| Orthologs & Meiotic GO | 78 | 6.5% | 253 | 21.0% | 164 | 13.6% | 387 | 32.1% | 324 | 26.9% |

Our analysis reveals that homeologous MGs on homeologs mostly show balanced expression and lack a significant change in MG content following polyploidization. The majority of homeologous genes (not only MGs) on homeologs also show over 95% sequence identity to each other [**37,74**]. Given these observations, such homeologs could synapse and recombine during meiosis. However, in allohexaploid wheat, homologs rather than homeologs synapse and recombine during meiosis ensuring the stability and fertility of this species, and the *Ph1* locus, in particular the *TaZIP4* gene copy inside this locus, has been identified as the main locus controlling this process. The wheat *ZIP4*, an ortholog of *ZIP4/Spo22* in *A. thaliana* and rice, is a member of the ZMM genes involved in the synaptonemal complex formation and class I crossovers pathway [**75,76**]. Moreover, wheat lacking *Ph1* exhibits extensive genome rearrangements, including translocation, duplications, and deletions [**33**]. Thus, the evolution of *Ph1* during wheat polyploidization may explain why wheat has largely maintained a similar gene content and balanced expression of its homeologs. How meiosis has adapted to cope with allopolyploidy in other species is still to be resolved; however, it has been suggested that reduction in the copy number of meiotic genes may stabilize the meiotic process after polyploidization [**35,66**]. The present study shows that this is not the case in wheat. It is possible that the presence of *Ph1* in wheat enabled the retention of multiple copies of meiotic genes as an alternative mechanism to ensure proper segregation of chromosomes during meiosis. In any case, the identification of the *TaZIP4* gene within the *Ph1* locus as the gene responsible for the *Ph1* effect on recombination and the observed effects of *Ph1* in wheat suggests that it may have more of a central role in meiosis than originally suspected from studies on model systems [**75,76**]. It has recently been suggested that *ZIP4* might acts as a scaffold protein facilitating physical interactions and assembly of different proteins complexes [**75,76,77**]. Therefore, our co-expression network was used to identify the wheat orthologs of known MGs connected with *TaZIP4.* The analysis indicates that the three *TaZIP4* homeologs on group 3 (TraesCS3A02G401700, TraesCS3B02G434600 and TraesCS3D02G396500) were clustered in module 2, the largest meiosis-related module, and strongly connected to many orthologs of MGs with various meiotic functions (**Fig 8**). However, the *TaZIP4* copy responsible for *Ph1* phenotype (TraesCS5B02G255100) did not cluster in the same module, reflecting its different expression profile from the other homeologs, being expressed in most tissues [**31-33**]. *TaZIP4* was connected to wheat orthologs of genes known to be involved in crossover formation such as *MSH2*, *SHOC1*, *FANCM*, *FLIP*, *EME1B*, *MUS81* [**2**]. This suggests that there may be an interplay between *TaZIP4* and genes from the anti-crossover pathway. The *TaZIP4* sub-network supports a more central role of *ZIP4* in meiosis than originally suspected from studies on model species.

**Fig 8.**
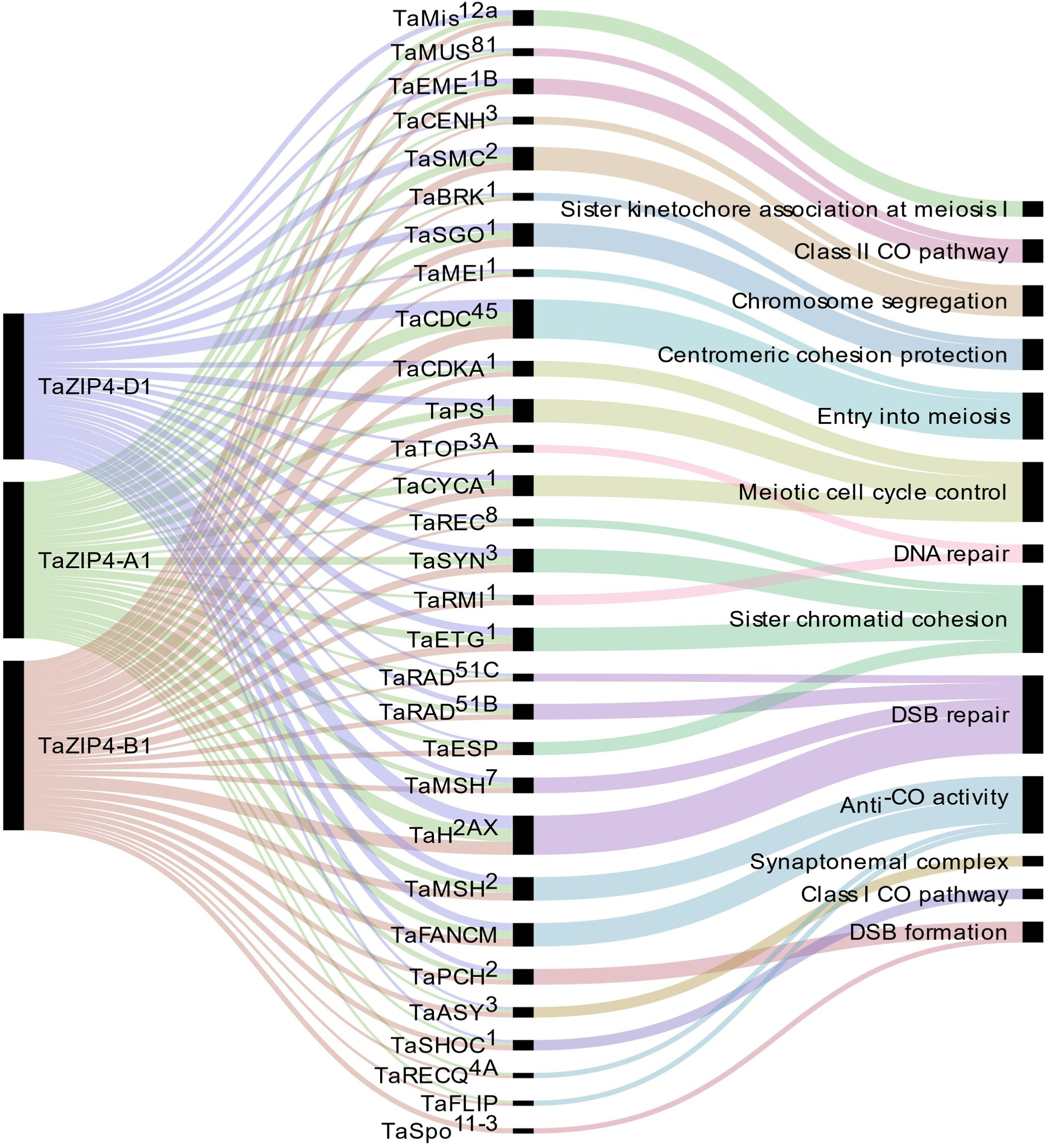
The wheat MG orthologs connected to *TaZIP4*. The alluvial diagram shows the connected genes to the *TaZIP4* homeologs *TaZIP4-A1*, *TaZIP4-B1* and *TaZIP4-D1*. Edge weight > 0.05 was used as threshold to visualise connected genes. Black bars indicate the number of homeologs for each connected gene.

### Further characterisation of the wheat meiotic co-expression network

#### 1) Identification of hub genes in the meiosis-related modules

Hub genes were identified within our meiosis-related modules by calculating the correlation between expression patterns of each gene and the module eigengene: the most highly correlated genes to the eigengene being the hub genes. The top 10 hub genes of each module with their functional annotation are shown in **Table 4**. The top 10 hub genes in module 2 were core histone genes, supporting the strong contribution of histones in this meiosis-related module. For further verification of histone involvement in module 2 and other modules in general, all wheat genes annotated as core histones or having GO terms related to histone modification, were retrieved for enrichment analysis. Analysis showed that the five types of histones (H1, H2A, H2B, H3 and H4) were enriched only in module 2 (*P* = 3.6 × 10^−4^, 1.2 × 10^−22^, 1.1 × 10^−19^, 9.4 × 10^−21^, and 3.3 × 10^−26^, respectively), having 433 genes (85% of all core histone genes in all modules), compared to an expected number of genes of 39 (**Fig 9**). Similar results were obtained for histone modification genes. Module 2 was the most enriched module with this group of genes (*P* = 9.3 × 10^−52^), containing 438 genes (30% of all histone modification genes in all modules). The histone modification genes were also enriched in 11 other modules, including the other meiosis-related modules (modules 28 and 41), however, with much lower numbers of enriched genes (**Fig 9**). Detailed information about genes included in this analysis is provided in **S14 Table**. The strong enrichment of histone modification genes in module 2 (the largest meiosis-related module) supports the important role of histone modifications in meiosis [**78-83**].

**Table 4.**
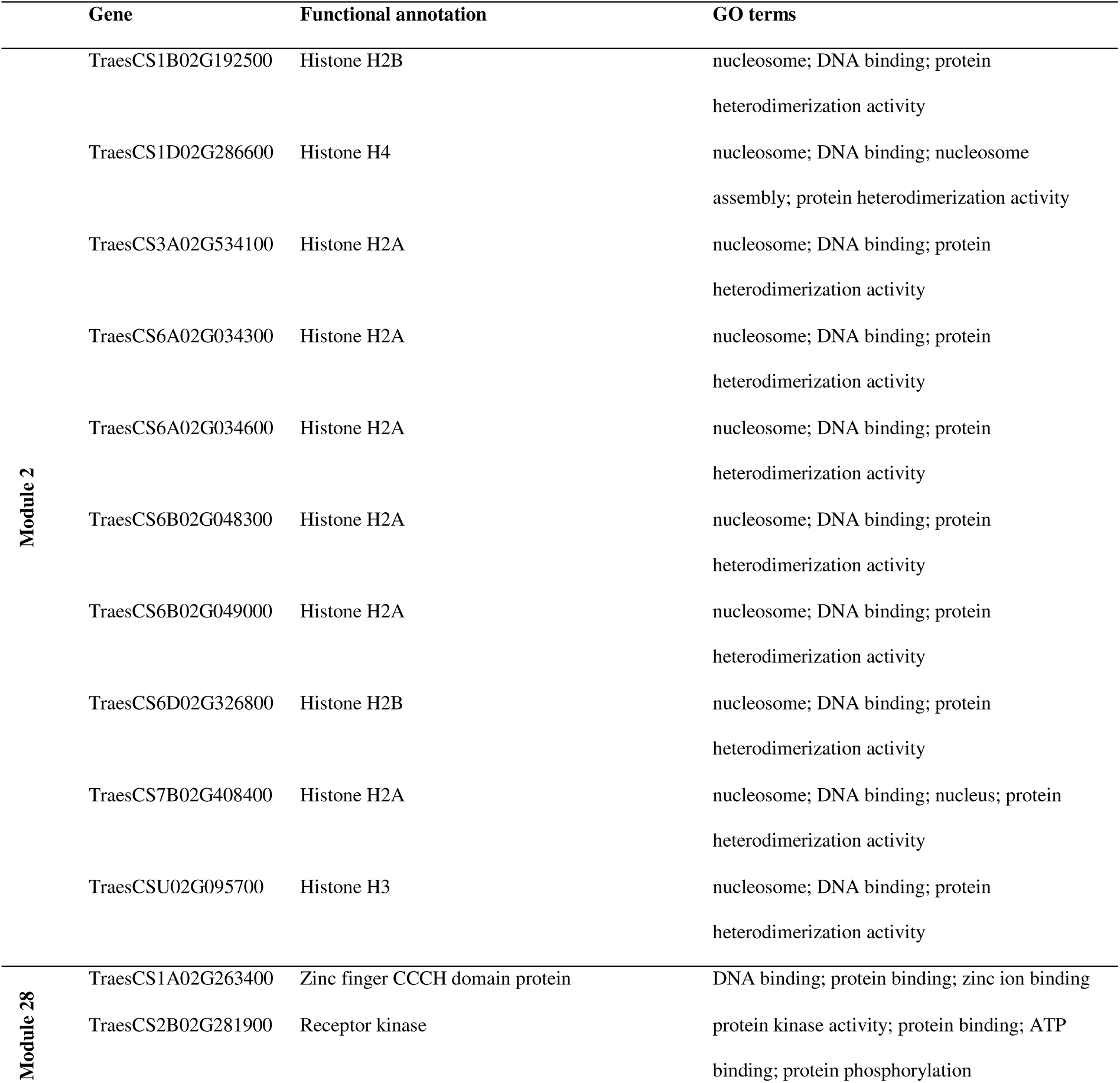
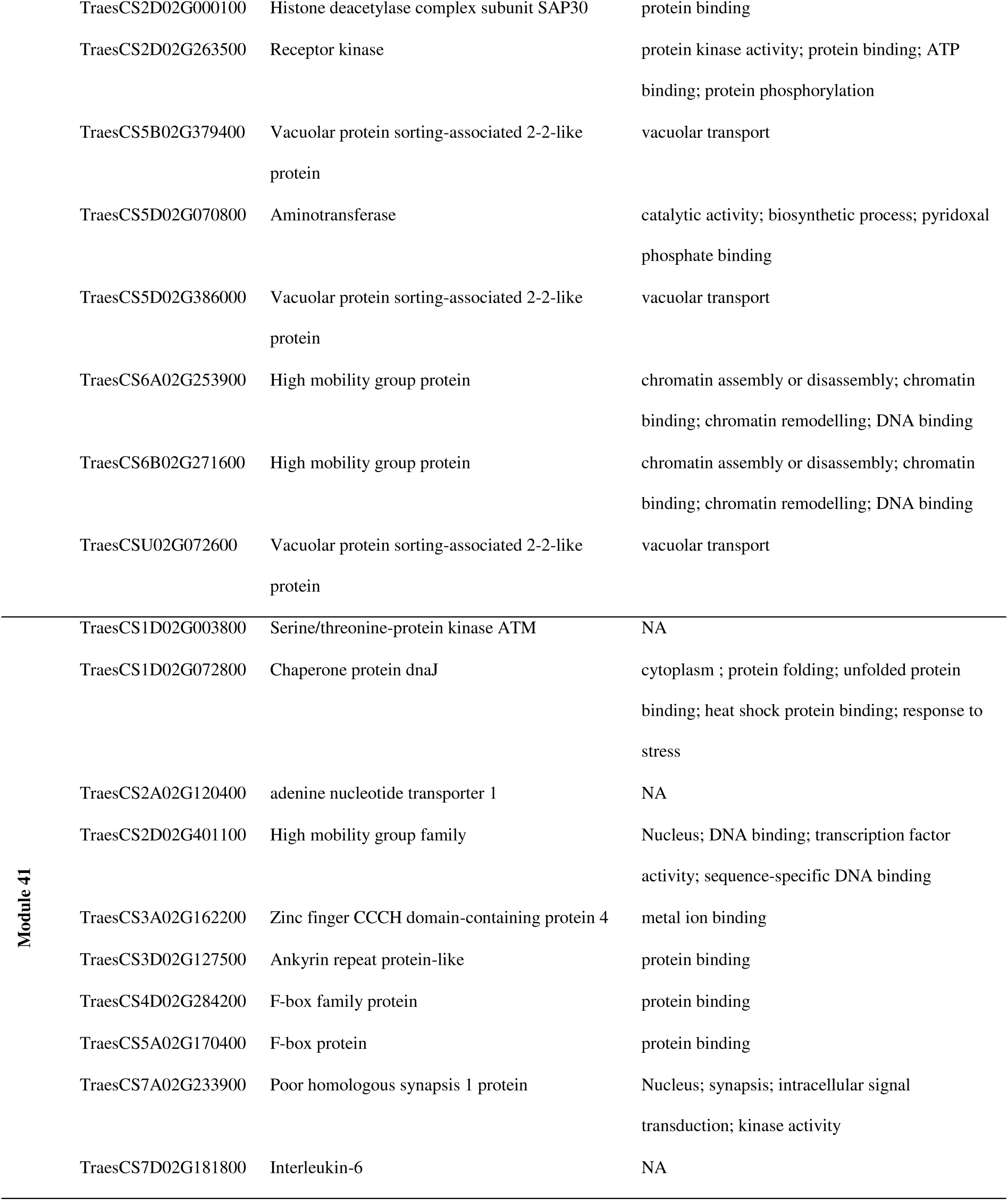
The top 10 hub genes of each meiosis-related module with their functional annotation.

**Fig 9.**
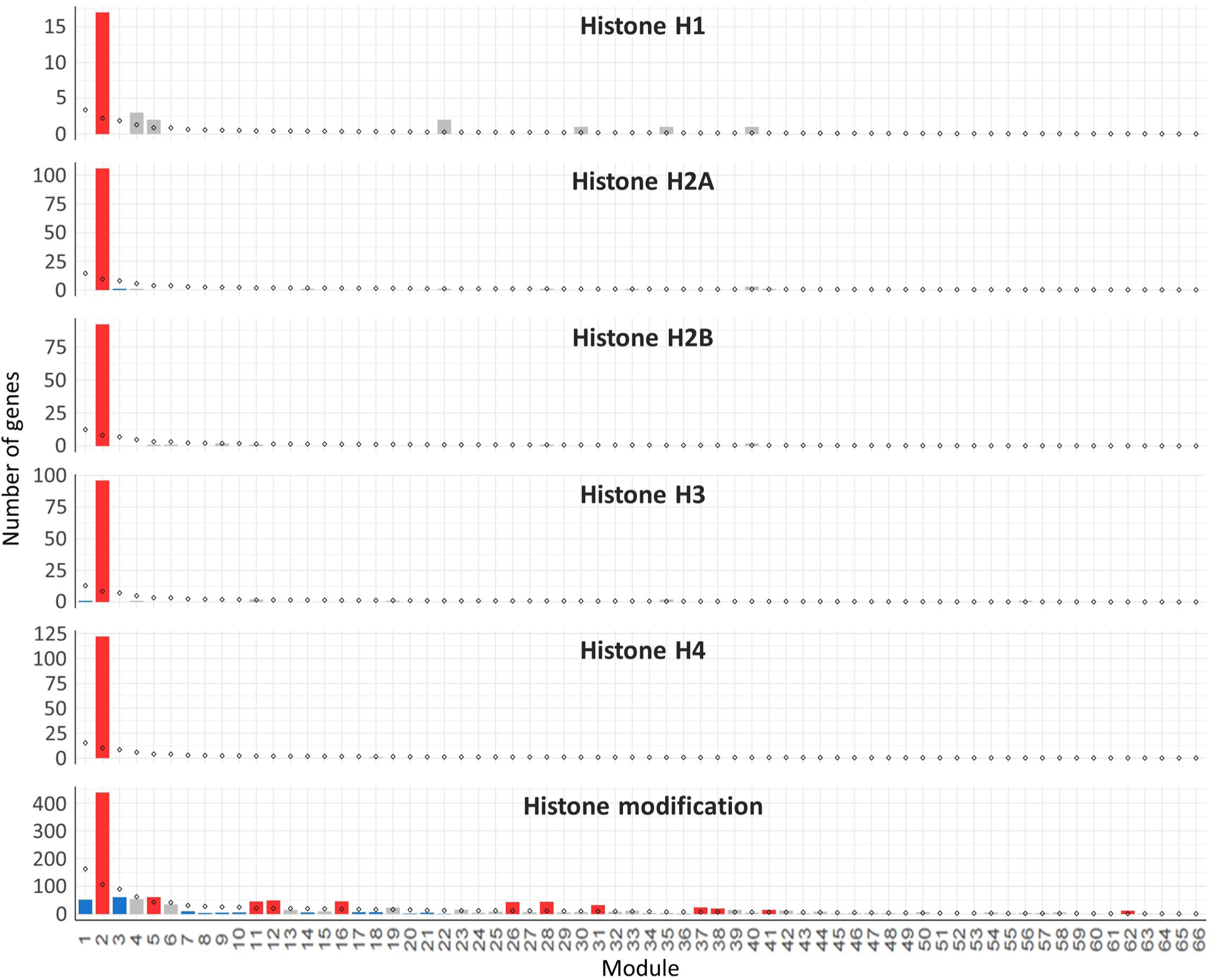
Histone genes enrichment in the gene co-expression network modules. The analysis included all the genes annotated as core histones (H1, H2A, H2B, H3 and H4) in the wheat genome and the genes with GO terms related to histone modification. Statistical significance of gene enrichment in a module at *P* < 0.05 is colour coded (Red indicates enriched, blue depleted and grey not significant). Rhombus shape indicates the expected number of genes in module.

Hub genes such as “*Poor homologous synapsis 1”* (*PHS1*) were also identified with module 41, the module most highly correlated to meiotic samples. This gene has been previously reported to have a key role in homologous chromosome pairing, synapsis, DNA recombination and accurate chromosome segregation during meiosis in maize [**84**], Arabidopsis [**85**] and wheat [**59**]. Other hub genes identified in modules 28 and 41 encoded for the high mobility group proteins [**86-90**], histone deacetylase [**91-94**], and F-box proteins [**95**].

#### 2) Analysis of Transcription factors within the meiosis-related modules

Many transcription factors (TFs) have been reported as key regulators of meiosis from studies on animals [**96,97**], yeast [**98-101**] and protozoa [**102**]. However, very little is known about the involvement of TFs in plant meiosis. The meiotic co-expression network was therefore exploited to identify potential meiosis-specific TFs. An assessment was undertaken of the enrichment of previously identified TF families in hexaploid wheat in the meiosis-related modules 2, 28 and 41. A total of 4889 high confidence genes belonging to 58 TF families were predicted from the annotation of the wheat genome sequence. Of these, 2439 TFs from 57 families could be assigned to the 66 modules in the gene co-expression network. Modules 2, 28 and 41 (meiosis-related modules) had 225, 25 and 17 TFs belonging to 31, 13 and 9 TF families, respectively (S15 Table). Compared to the expected number of TF families genes in each module, only 5 TF families were significantly enriched in module 2: Mitochondrial transcription termination factor (mTERF); Growth-Regulating Factor (GRF); abscisic acid-insensitive protein 3/Viviparous1 (ABI3/VP1); Forkhead-associated domain (FHA) and E2F/ Dimerization Partner (DP). On the other hand, 4 TF families were significantly depleted (**Fig 10**). The TF family NAC was the only TF family significantly enriched in module 41, containing 5 NAC genes (expected number 0.6; *P* < 0.05). Module 28 was not enriched with any TF family, although E2F/DP transcription factors were enriched in this module with borderline statistical significance (*P* = 0.06), with 4 genes in this module (the expected number was 0.2). Except in module 2, E2F/DP and FHA transcription factors families were not enriched in any other modules in the gene co-expression network (**S16 Table**). E2F/DP plays an important role in regulating gene expression necessary for passage through the cell cycle in mammals and plants [**103-105**]. Members of FHA contain the forkhead-associated domain, a phosphopeptide recognition domain found in many regulatory proteins. Genes belonging to the FHA group are reported to have roles in cell cycle regulation [**106-108**], DNA repair [**109-112**] and meiotic recombination and chromosome segregation [**113-115**]. A previous meiotic transcriptome study identified up-regulation of TFs belonging to the MADS-box, bHLH, bZIP, and NAC families in Arabidopsis and maize meiocytes at early meiosis [**116**]. Zinc finger-like TFs have also been suggested to be regulators of maize MG expression [**117**]. The present study indicates that TF families known to have roles in cell cycle and meiosis processes are over-represented in the meiosis-related modules (module 2 in particularly). Those TF families contain about 20 meiosis-specific candidate TF genes whose function can be validated using the available reverse genetics resources in polyploid wheat [**118**].

**Fig 10.**
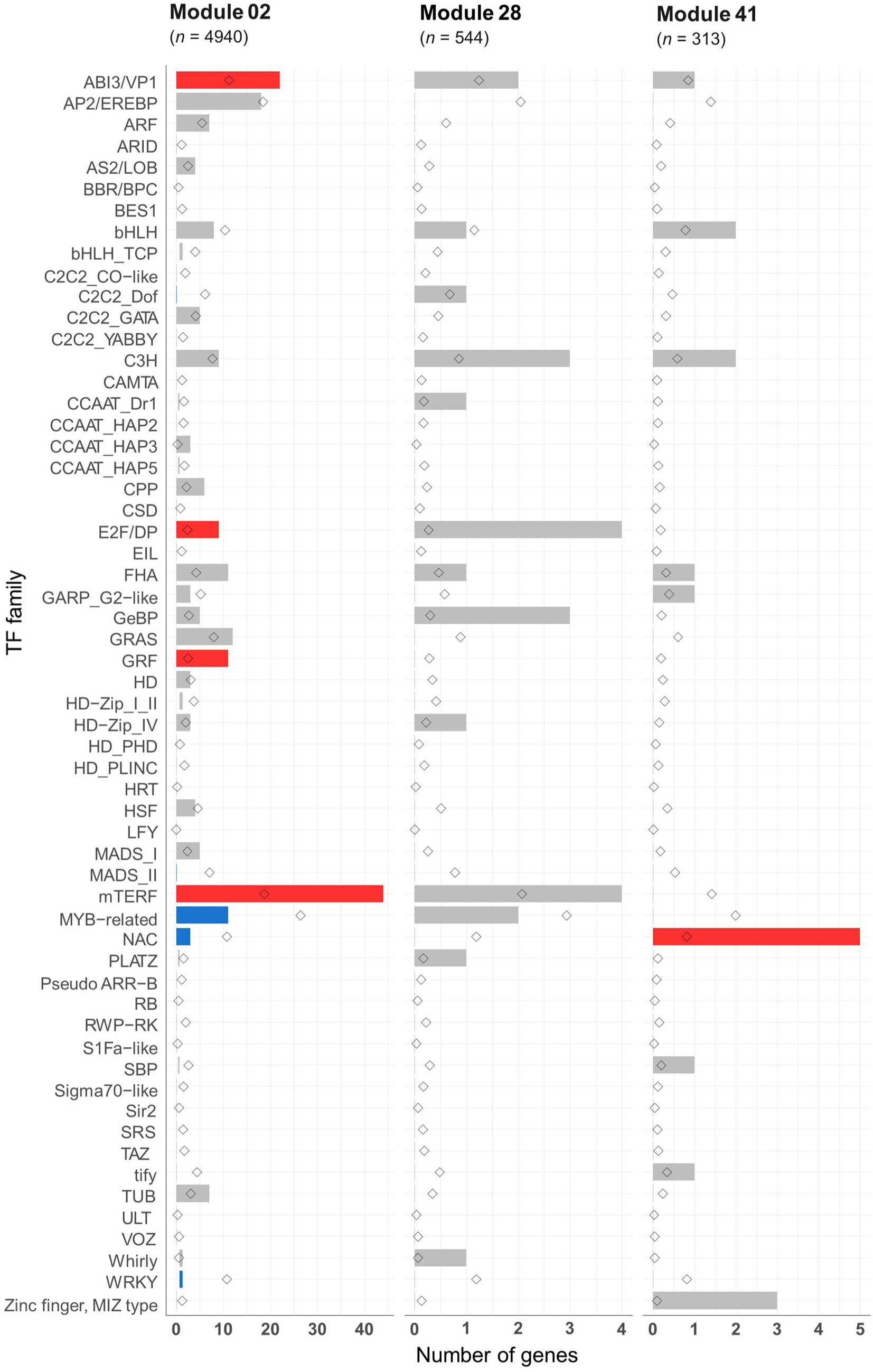
Transcription factor families in the meiosis-related modules. Statistical significance of gene enrichment in the modules is colour coded (Red indicates over-represented, blue under-represented and grey not significant; *P* < 0.05). Rhombus shape indicates the expected number of genes in module.

#### 3) Visualisation of networks and identification of candidate MGs

Having identified meiosis-related modules, the Networks within such modules can be visualised, highlighting genes for future studies. Edge files were created with gene annotation for the three meiosis-related modules 2, 28 and 41. Those files can be used to investigate the relation between orthologs of MGs within a module and ranked based on the strength of the connection (weight value). Another application of co-expression networks is the identification of previously uncharacterised genes regulating biological processes [**37,40,42-49**]. Cytoscape 3.7.1 software [**119**] was therefore used to visualise our network and to show connections between different orthologs of MGs in meiosis-related modules. Wheat MG orthologs in meiosis-related modules were used as “guide genes” to construct co-expression subnetworks containing only genes with direct connections to the guide genes. One such subnetwork is shown in **Fig 11**, where the following wheat orthologs of MGs in module 41 were selected and used to construct a meiotic subnetwork: *Poor Homologous Synapsis 1* (*TaPHS1;* [**59**]); *Argonaute* (AGO9/AGO104; [**120,121**] *Replication protein A2c* (*OsRPA2c*; [**122**]; *Meiotic nuclear division protein 1* (*AtMND1*; [**123,124**]); *MMS and UV Sensitive 81* (*AtMUS81*; [**125,126**]); and *Parting Dancers* (*AtPTD*; [**127,128**]) (guide genes; red circles in **Fig 11**). The network complexity was reduced using an edge weight > 0.05. The visualized subnetwork contained 53 gene IDs including 9 guide gene copies. The gene TraesCS7A02G233900 (*TaPHS1*), a hub gene in module 41, was central in the network having the highest number of direct edges (41 direct edges; connected with 77.4% of the genes in the subnetwork). This subnetwork allowed identification of other genes with putative roles in meiosis (pink circles): a) RNA recognition motifs-containing gene (TraesCS5A02G319000) similar to Mei2, a master regulator of meiosis and required for premeiotic DNA synthesis as well as entry into meiosis in *Schizosaccharomyces pombe* [**129,130**]; b) the gene TraesCS4D02G050000 showed similarity to Male meiocyte death 1 (*MMD/DUET*), a PHD-finger protein plays role in chromatin structure and male meiotic progression in *A. thaliana* [**131,131**]; c) the gene TraesCS5D02G454900, a possible TF belonging to the FHA family known to have function in cell cycle regulation [**106-108**], DNA repair [**109-112**] and meiotic recombination and chromosome segregation [**113-115**]. The meiotic subnetwork contained genes with similarity to cell cycle like F-box family proteins, high mobility family proteins, and chromatin remodelling genes. The subnetwork also contained a group of genes connected to most of our guide genes, which thus might be involved with them in similar biological processes. Examples of such genes are TraesCS3A02G101000, TraesCS1A02G292700 and TraesCS1D02G291100 which encode for zinc finger CCCH domain-containing proteins (**Fig 11**). Other meiotic subnetworks were also constructed using other guide genes from modules 2 and 28.

**Fig 11.**
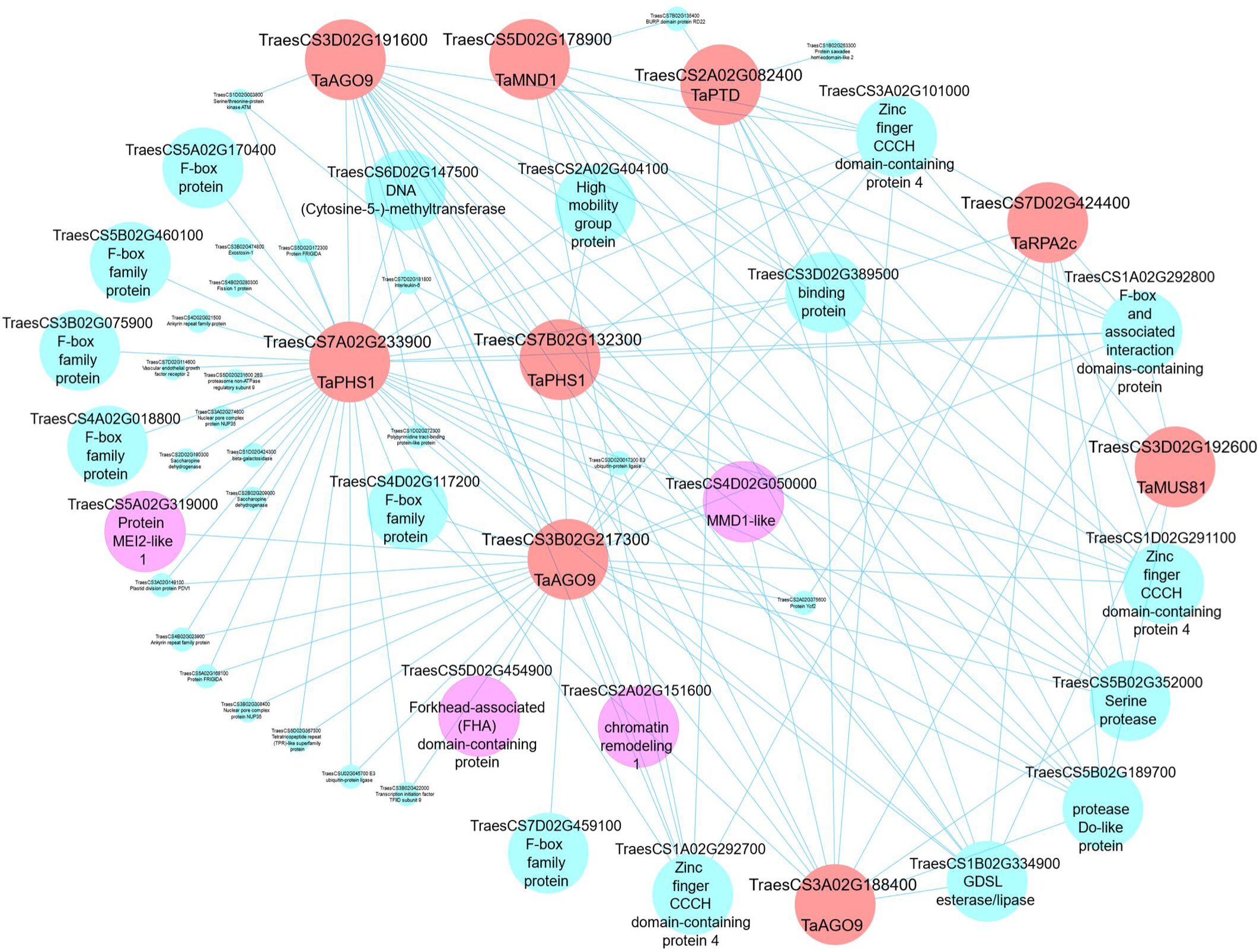
A meiotic co-expression subnetwork in hexaploid wheat. This subnetwork was constructed using 9 guide genes in module 41. Guide genes are wheat orthologs of MGs in other plant species (red circles); pink circles represent genes with putative meiosis function. Edge weight 0.05 was used as threshold to visualise genes in the subnetwork using Cytoscape 3.7.1 software.

#### 4) The meiotic co-expression network is accessible in a larger biological contest

Our WGCNA co-expression network and GO enrichment data has been integrated with the wheat knowledge network [**132**] to make it publicly accessible in a larger biological context and to make it searchable through the KnetMiner web application (http://knetminer.rothamsted.ac.uk; [**133**]. KnetMiner can be searched with keywords (incl. module ID and GO terms) and wheat gene identifiers. The gene knowledge graphs generated contain many additional relation types such protein-protein interactions, homology and links to genome wide association studies and associated literature placing the co-expression networks generated here in a wider context.

## Conclusion

In summary, the present study shows that most MGs in wheat are retained as three homeologous genes, which are expressed during meiosis at similar levels, suggesting that they have not undergone extensive gene loss nor sub/neo-functionalisation. Meiosis-related modules have been used to create networks and identify hub genes providing targets for future studies. The network containing the *ZIP4* gene, recently defined as *Ph1* [**31-33**] for example, highlights potential interacting partners. Finally, the networks highlight genes such as *ZIP4* and “*Poor homologous synapsis 1*”, which may play a more central role in meiosis than previously thought. The co-expression network analysis combined with orthologue information will contribute to the discovery of new MGs and greatly empowers reverse genetics approaches to validate the function of candidate genes [**118**]. Ultimately this will lead to better understanding of the regulation of meiosis in wheat (and other polyploid plants) and subsequently improve wheat fertility.

## Materials and Methods

### RNA-Seq data collection

For co-expression network analysis we included 130 samples, containing 113 samples previously described in Ramírez-González et al. [**36**] and 17 samples from anthers during meiosis (9 samples from Martin *et al*. [**33**]), and 8 samples downloaded from (https://urgi.versailles.inra.fr/files/RNASeqWheat/Meiosis/). Samples were selected to represent all main tissue types: grain (*n* = 37 samples), leaves (*n* = 21 samples), roots (*n* = 20 samples), anther at meiosis (*n* = 17 samples), spike (*n* = 12 samples), floral organs (anther, pistil and microspores) at stages other than meiosis (*n* = 10 samples), stem (*n* = 7 samples) and shoots (*n* = 6 samples). All samples were under nonstress conditions and mostly from the reference accession Chinese Spring. Detailed information about the used samples are listed in the supplementary materials (**S2 Table**).

### Mapping of RNA-Seq reads to reference

Kallisto v0.42.3 [**134**] was used to map all RNA-Seq samples to the Chinese Spring transcriptome reference IWGSC RefSeq Annotation v1.1 [**37**], following default parameters previously shown to result in accurate homeolog-specific read mapping in polyploid wheat [**36,50**]. Tximport v1.2.0 was then used to summarise expression levels from transcript to gene level **(S1 Text)**.

### Co-expression network construction

The WGCNA package in R [**51,52**] was used to construct the scale-free co-expression network. Metadata for all samples were assigned with 8 tissue types (average 16.25; median 14.5 replicates per factor). Only high confidence (HC) genes [**37**] with expression > 0.5 TPM in at least one meiosis sample were retained for co-expression network construction using the R Package WGCNA (version 1.66) [**51,52**]. Using the varianceStabilizingTransformation() function from DESeq2 [**135**], the count expression level of selected genes was normalised to eliminate differences in sequencing depth between different RNA-Seq studies **(S2 Text)**. To select a soft power threshold (β) for adjacency calculation (as a*_ij_* = |s*_ij_*|^β^; where s*_ij_* is the correlation between gene *i* and gene *j*), the Scale-free Topology Criterion was used [**136**]. Using the pickSoftThreshold() function to calculate β values, the soft power threshold emphasising strong correlations between genes and penalising weak correlations was selected as the first power to exceed a scale-free topology fit index of 0.9 [**36**] **(S3 Text)**. The correlation type used to calculate adjacency matrices was biweight midcorrelation (bicor). The adjacency matrices were transformed into a topological overlap matrix (TOM), measuring the network connectivity of a gene defined as the sum of its adjacency with all other genes for network generation. The blockwiseModules() function was used to calculate matrices and construct blockwise networks considering the following parameters: network type (networkType) = “signed hybrid”, maximum block size (maxBlockSize) = 46,000 genes, soft power threshold (power) = 7, correlation type (corType) = “bicor” (biweight midcorrelation with maxPOutliers set to 0.05 to eliminate effects of outlier samples), topological overlap matrices type (TOMType) = “unsigned” with the mergeCutHeight = 0.15 and the minModuleSize = 30 to classify genes with similar expression profiles into gene modules using average linkage hierarchical clustering, according to the TOM-based dissimilarity measure with a minimum module size of 30 genes **(S4 Text)**. Module Eigengene (MEs), summarising the expression patterns of all genes within a given module into a single characteristic expression profile, were calculated as the first principal component in the Principal Component Analysis (PCA) using the moduleEigengenes() function **(S4 Text)**.

### Identifying meiosis-related modules

The module eigengenes was used to test correlations between gene modules and traits (8 tissue types) using the cor() function. To assess the significance of correlations, Student asymptotic *P* values for correlations were calculated using the function corPvalueStudent(), and corrected for multiple testing by calculating FDR (false discovery rate) using a p.adjust() function following the Benjamini & Yekutieli [**137**] method. We considered a module meiosis-related when its correlation was strong with meiosis samples (*r* > 0.5 and FDR < 0.05) and weak (*r* < 0.3) or negative with other type of tissues **(S5 Text)**.

### Analysis of GO term enrichment in modules

GO term enrichment was calculated using the “goseq” package [**138**]. Gene ontology (GO) annotations of IWGSC RefSeq v1.0 genes were retrieved from the file “FunctionalAnnotation.rds” in https://opendata.earlham.ac.uk/wheat/under_license/toronto/Ramirez-Gonzalez_etal_2018-06025-Transcriptome-Landscape/data/TablesForExploration/FunctionalAnnotation.rds [**36**] by filtering for ontology “IWGSC+Stress”. GO data was then converted to IWGSC RefSeq Annotation v1.1 by replacing “01G” by “02G” in the IWGSC v1.0 gene IDs and retaining only genes > 99 % similar with > 90% coverage in the v1.0 and v1.1 annotation versions (as determined by blastn of the cDNAs) (called “all_go”). *P* values for GO term enrichment were calculated using the goseq() function (using the following parameters: the pwf object was created using the nullp() function which calculated a Probability Weighting Function for the genes v1.1 based on their length, the gene2cat = all_go, and test.cats = “GO:BP”, to specify the Biological Process GO term category to test for over representation amongst the inquired genes) and corrected using the FDR method [**139**]. A GO term was considered enriched in a module when FDR adjusted *P* value < 0.05 **(S6 Text)**. All figures shown for enriched GO terms in the modules were produced using RAWGraphs software [**140**].

### Orthologs of MGs in wheat

A comprehensive literature search was performed for MGs in model plant species (mainly *A. thaliana* and rice; **S17 Table**), identifying gene IDs based on the “Os-Nipponbare-Reference-IRGSP-1.0” for rice (*Oryza sativa* Japonica Group) and “TAIR10” for *A. thaliana*. Wheat orthologs of MGs were then retrieved from *Ensembl* Plants Genes 43 database through BioMart (in; http://plants.ensembl.org/biomart) where orthologs calculated according to Vilella et al. [**141**] using the following gene datasets: “Triticum aestivum genes (IWGSC)”, “Oryza sativa Japonica Group genes (IRGSP-1.0)” and “Arabidopsis thaliana genes (TAIR10)” for wheat [**37**], rice [**142,143**] and Arabidopsis [**144**], respectively. For genes with no orthologs identified using this method, potential wheat orthologs were identified by searching for amino acid sequence similarity using BLASTP [**145**] in *Ensembl*Plants according to the following criteria: E-value < 1e-10; ID% > 25% with Arabidopsis and > 70% with rice; then only wheat genes for which *Ensembl*Plants did not list any orthologs in rice and Arabidopsis were considered as orthologs of MGs. Finally, 407 wheat gene IDs were identified as orthologs of 103 plant MGs (listed in **S5 Table**). This group of genes was referred to in this study as “Orthologs”.

### Wheat genes with MG ontology (GO)

A total number of 46,909 GO terms used by Ramírez-González et al. [**36**] to calculate GO term accessions for wheat genes (IWGSC v1.0 gene annotation) were filtered for meiosis-related GO terms, using 15 meiosis-specific keywords (“meiosis”, “meiotic”, “synapsis”, “synaptonemal”, “prophase I”, “metaphase I”, “anaphase I”, “telophase I”, “leptotene”, “zygotene”, “pachytene”, “diplotene”, “chiasma”, “crossover” and “homologous chromosome segregation”). A total of 284 meiosis GO accessions were identified and used to retrieve 927 wheat genes with potential roles during meiosis (**S7 Table**). All genes identified by gene orthologs and gene ontology methods were then filtered to retain only genes had expression > 0.5 TPM in at least one meiosis sample. This group of genes was referred to in this paper as “Meiotic GO”. Enrichment analysis for the genes from “Orthologs” and “Meiotic GO” groups in all module was conducted (**S7 Text**). The number of genes from each group was assessed in all modules and compared with the expected number based on the module size. There was a set of genes overlapping between “Orthologs” and “Meiotic GO” groups, which was considered in the “Orthologs” group when undertaking gene enrichment analysis. Fisher’s exact test was used to calculate significant enrichment in the modules. Gene group considered over- or under-represented in a module when *P* < 0.5.

### Identifying highly connected hub genes

Hub genes within each module were identified using the WGCNA R package function signedKME() to calculate the correlation between expression patterns of each gene and the module eigengene. Hub genes were considered those more highly correlated to the eigengene **(S8 Text)**.

### Assessment of TF families in modules

A total of 4889 wheat HC genes (IWGSC RefSeq Annotation v1.1; [**37**]) belonging to 58 TF families were predicted from the annotation of the wheat genome sequence (https://github.com/Borrill-Lab/WheatFlagLeafSenescence/blob/master/data/TFs_v1.1.csv). The number of TFs from each family was assessed in all modules and compared with the expected number based on the module size. Fisher’s exact test was used to calculate significant enrichment of TFs in the modules. TF family considered over- or under-represented in a module when *P* < 0.5 **(S9 Text)**.

### Defining gene categories based on number of homeologs

A list of homeologs for all HC hexaploid wheat genes (IWGSC v1.1 gene annotation; [**37**]) was retrieved from *Ensembl* Plants Genes 43 database through BioMart (in; http://plants.ensembl.org/biomart). Based on number of homeologs from each of the A-, B- and D-sub-genomes, genes were assigned to four groups: triads that refer to 1:1:1 triads (with a single copy from each of the A-, B- and D-sub-genomes); duplets referring to 1:1:0, 1:0:1 and 0:1:1 duplets; monads group containing genes with no homeologs (e.g. 0:0:1); and “others” containing genes with more than two homeologs, in conjunction with genes from the homeologous groups 0:1:2, 0:2:1, 1:0:2, 2:0:1, 1:2:0, 2:1:0, 2:0:0, 0:2:0 and 0:0:2. Accordingly, 19801 triads (59403 genes), 7565 duplets (15130 genes), 15109 monads (single-copy genes) and 18250 genes from the “Others” group were identified (**S1 Table**).

### Defining gene categories based on homeolog expression patterns in triads

Homeolog expression pattern in triads was determined for each of the eight tissue types (**S10 Text**, first part). For triads it was calculated according to Ramírez-González *et al*. [**36**] where a triad can be described as balanced, A dominant, A suppressed, B dominant, B suppressed, D dominant or D suppressed, based on the relative expression contribution of its A, B and D homeologs. Triads were defined as expressed when one of its homeologs was expressed according to the criterion used in our WGCNA analysis (**S18 Table**). This insured that all triads contain genes from modules were included in the homeolog expression bias analysis (**S10 Text**, second part). Genes from a triad might not belong to the same module due to dissimilarity of their expression patterns. Thus, to allow the assessment of the expression pattern of genes in each module, each homeolog (A, B and D homeologs) in a triad was assigned to one of the three categories “Balanced”, “Dominant” and “Suppressed” based on the homeolog origin (A, B and D sub-genome) and the triad description (balanced, A dominant, A suppressed, B dominant, B suppressed, D dominant or D suppressed) as shown in **S19 Table**. The values of the relative contributions of each homeolog per triad were used to plot the ternary diagrams using the R package ggtern [**146**].

### Co-expression gene network visualisation

Cytoscape software (version 3.7.1; [**119**]) was used to visualise the network described in this study. Firstly, the “exportNetworkToCytoscape” function was used to create edge files which could be used to visualize the network, then depending on network complexity, different weight value thresholds were used to filter genes to be visualised (**S11 Text**,). The term ‘weight value’ in the input files for Cytoscape refers to the connection strength between two nodes (genes) in terms of correlation value obtained from the topological overlap matrices (TOM). The co-expression network data has also been integrated with the wheat knowledge network [**132**] to make it publicly accessible through the KnetMiner web application (http://knetminer.rothamsted.ac.uk; [**133**]. The data was semantically modelled as nodes of type Gene, Co-Expression-Module, Co-Expression-Study, GOterm; connected by relations of type part-of and enriched. Each module was given a unique identifier composed of the module number and the prefix “AKA”. KnetMiner can be searched with keywords (incl. module ID and GO terms) and wheat gene identifiers.

## Data availability

All data files used in this manuscript are deposited in the Earlham Institute Open Data Platform (https://opendata.earlham.ac.uk/wheat/under_license/toronto/Martin_etal_2018_Alabdullah_etal_2019_wheat_meiosis_transcriptome_and_co-expression_network/). All R scripts are provided as .text files in the Supporting Information. The Gene network data will become accessible and searchable through the public plant website, KnetMiner (http://knetminer.rothamsted.ac.uk) with its next update this August.

## Supporting information

S1 Table. Assignment of hexaploid wheat HC genes to four groups based on their copy number

S1 Text. R script used to summarize expression values (counts) from transcript to gene level

S2 Table. The 130 RNA-Seq samples included in the co-expression analysis

S2 Text. R script used to combine samples from all studies and normalize counts for WGCNA analysis

S3 Table. Number of genes and gene IDs in the modules identified using WGCNA

S3 Text. R script used to calculate the soft power threshold for the WGCNA analysis

S4 Table. Enriched GO terms in all modules including the three meiosis-related modules

S4. Text. R script used to run the WGCNA analysis and MEs calculation and plotting

S5 Table. Gene IDs of wheat orthologs of known meiotic genes in other plants species

S5. Text. R scripts used to calculate module-tissue correlation

S6 Table. Meiotic genes and their orthologs in wheat according to EnsemblPlants prediction

S6. Text. R scripts used for enrichment analysis of GO and GO slim terms in the modules

S7 Table. List of wheat genes with meiotic GO term(s)

S7. Text. R scripts used for enrichment analysis of ???????Orthologs??????? and ???????Meiotic GO??????? genes in the modules

S8 Table. Fishers exact test and FDR adjusted p-value for "Orthologs" and "Meiotic GO" genes enrichment in modules

S8 Text: R script used to calculate hub genes for each module

S9 Table. Assignment of wheat orthologs of known meiotic genes to the meiosis-related modules

S9. Text. R script used for the assessment of TF families in modules

S10 Table. Expression pattern of wheat orthologs of meiotic genes that were not assigned to the meiosis-related modules

S10 Text. R script used to calculate homeolog expression patterns in triads

S11 Table. Copy number of orthologs of meiotic gene in wheat and its ancestors

S11. Text. R script used to export meiosis-related modules data for visualisation

S12 Table. Assignment of the genes in triads to the co-expression network modules

S13 Table. HC genes distribution across wheat genomic compartment

S14 Table. List of the core histone and histone modification genes identified in the wheat genome

S15 Table. List of transcription factors in the meiosis-related modules 2, 28 and 4

S16 Table. Fishers exact test for TF families enrichment in modules

S17 Table. Literature review of the functionally characterized meiotic genes in model plant species

S18 Table. Defining the expressed triads in meiotic anther tissue

S19 Table. Gene category assignment based on the sub-genome of origin and triad description

S1 Fig. Homeolog expression patterns of expressed triads in hexaploid wheat

S2 Fig. Proportion of genes in each homeologs number category

S3 Fig. Module-tissue relationship

S4 Fig. Enriched GO terms in the meiosis-related and other tissue-related modules

## Acknowledgements

This work was supported by the UK biotechnology and Biological Sciences Research Council (BBSRC) through a grant as part of Designing Future Wheat (DFW) Institute Strategic Programme (BB/P016855/1) and response mode grant (BB/R007233/1).

## Author Contributions

**Conceptualization:** Abdul Kader Alabdullah, Philippa Borrill, Azahara C. Martin, Cristobal Uauy, Graham Moore.

**Data curation:** Abdul Kader Alabdullah, Azahara C. Martin, Ricardo H. Ramirez-Gonzalez, Keywan Hassani-Pak.

**Formal analysis:** Abdul Kader Alabdullah

**Funding acquisition:** Graham Moore, Peter Shaw.

**Methodology:** Abdul Kader Alabdullah, Philippa Borrill, Cristobal Uauy, Graham Moore.

**Software:** Abdul Kader Alabdullah, Philippa Borrill, Ricardo H. Ramirez-Gonzalez, Keywan Hassani-Pak.

**Supervision:** Graham Moore, Peter Shaw.

**Writing – original draft:** Abdul Kader Alabdullah

**Writing – review & editing:** Abdul Kader Alabdullah, Philippa Borrill, Azahara C. Martin, Ricardo H. Ramirez-Gonzalez, Keywan Hassani-Pak, Cristobal Uauy, Peter Shaw and Graham Moore.

## Supporting information

**S1 Fig. Homeolog expression patterns of expressed triads in hexaploid wheat.** Homeolog expression pattern was calculated for 19,801 triads (59,403 genes) across 8 tissue types according to published criteria **[36]**, where triad defined as expressed when the sum of the A, B, and D subgenome homoeologs was > 0.5 TPM. **(A)** Proportion of triads in each homoeolog expression pattern across the 8 tissues. *n* is number of expressed triads. **(B)** Ternary plot showing relative expression abundance of 14,837 expressed triads (44,511 genes) in the meiotic anther tissue. Each circle represents a gene triad with an A, B, and D coordinate consisting of the relative contribution of each homeolog to the overall triad expression. Triads in vertices correspond to single-subgenome–dominant categories, whereas triads close to edges and between vertices correspond to suppressed categories. Box plots indicate the relative contribution of each subgenome based on triad assignment to the seven categories. Percentages between brackets indicate the percentage of triad number in each category to the total number of triads.

**S2 Fig. Proportion of genes in each homeologs number category.** Expressed genes across 8 tissues were assigned to four categories (triads, duplets, monads and others). *n* indicates number of expressed genes.

**S3 Fig. Module-tissue relationship.** Each row corresponds to a module; each column corresponds to a tissue type; Each cell contains the correlation value (*r*) and, in brackets, its corresponding FDR adjusted *P* value. *n* indicates number of samples. Only modules that have correlation value > 0.5 are shown.

**S4 Fig. Enriched GO terms in the meiosis-related and other tissue-related modules.** Top 5 GO terms are shown for each module. Black bars indicate the number of genes in the GO term.

**S1 Table. Assignment of hexaploid wheat HC genes to four groups based on their copy number.** *n* is number of samples per tissue type.

**S2 Table. The 130 RNA-Seq samples included in the co-expression analysis.** The samples belong to eight different types of tissues (Intermed.tissue).

**S3 Table. Number of genes and gene IDs in the modules identified using WGCNA.**

**S4 Table. Enriched GO terms in all modules including the three meiosis-related modules.**

**S5 Table. Gene IDs of wheat orthologs of known meiotic genes in other plants species.**

**S6 Table. Meiotic genes and their orthologs in wheat according to EnsemblPlants prediction.**

**S7 Table. List of wheat genes with meiotic GO term(s)**

**S8 Table. Fishers exact test and FDR adjusted p-value for “Orthologs” and “Meiotic GO” genes enrichment in modules.**

**S9 Table. Assignment of wheat orthologs of known meiotic genes to the meiosis-related modules.**

**S10 Table. Expression pattern of wheat orthologs of meiotic genes that were not assigned to the meiosis-related modules.** Average expression values (TPMs) is shown for the 8 tissue types.

**S11 Table. Copy number of orthologs of meiotic gene in wheat and its ancestors.** MR indicates wheat orthologs of known meiotic recombination genes. ZMM indicates orthologs of ZMM pathway genes.

**S12 Table. Assignment of the genes in triads to the co-expression network modules.** Genes belong to Meiotic, Non-meiotic, Leaves, Grain and Roots modules and “Orthologs & Meiotic GO” gene sets are indicated in the table. Balance as a gene category refer to the genes belong to balanced triads.

**S13 Table. HC genes distribution across wheat genomic compartment.**

**S14 Table. List of the core histone and histone modification genes identified in the wheat genome.** All wheat genes annotated as core histones (Fun.annotation) or having GO terms related to histone modification (GO) were selected.

**S15 Table. List of transcription factors in the meiosis-related modules 2, 28 and 4.**

**S16 Table. Fishers exact test for TF families enrichment in modules.**

**S17 Table. Literature review of the functionally characterized meiotic genes in model plant species.**

**S18 Table. Defining the expressed triads in meiotic anther tissue.** Triad is considered as expressed when one of its homeologs is expressed according to the criterion used in our WGCNA analysis.

**S19 Table. Gene category assignment based on the sub-genome of origin and triad description.**

**S1 Text. R script used to summarize expression values (counts) from transcript to gene level.**

**S2 Text. R script used to combine samples from all studies and normalize counts for WGCNA analysis.**

**S3 Text. R script used to calculate the soft power threshold for the WGCNA analysis.**

**S4. Text. R script used to run the WGCNA analysis and MEs calculation and plotting.**

**S5. Text. R scripts used to calculate module-tissue correlation.**

**S6. Text. R scripts used for enrichment analysis of GO and GO slim terms in the modules.**

**S7. Text. R scripts used for enrichment analysis of “Orthologs” and “Meiotic GO” genes in the modules.**

**S8 Text: R script used to calculate hub genes for each module.** Top 10 hub genes were selected for each module.

**S9. Text. R script used for the assessment of TF families in modules.**

**S10 Text. R script used to calculate homeolog expression patterns in triads.**

**S11. Text. R script used to export meiosis-related modules data for visualisation.**

