## Supplementary figures and images for "A co-expression network in hexaploid wheat reveals mostly balanced expression and lack of significant gene loss of homeologous meiotic genes upon polyploidization"

### S1 Fig. Homeolog expression patterns of expressed triads in hexaploid wheat

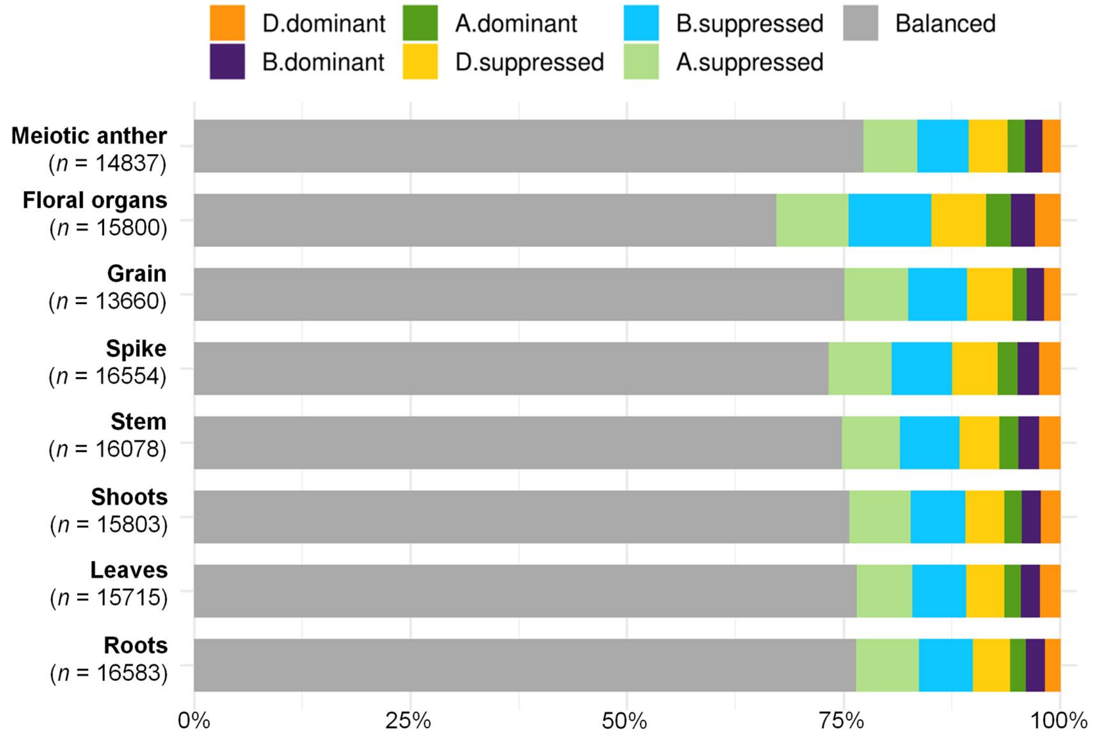

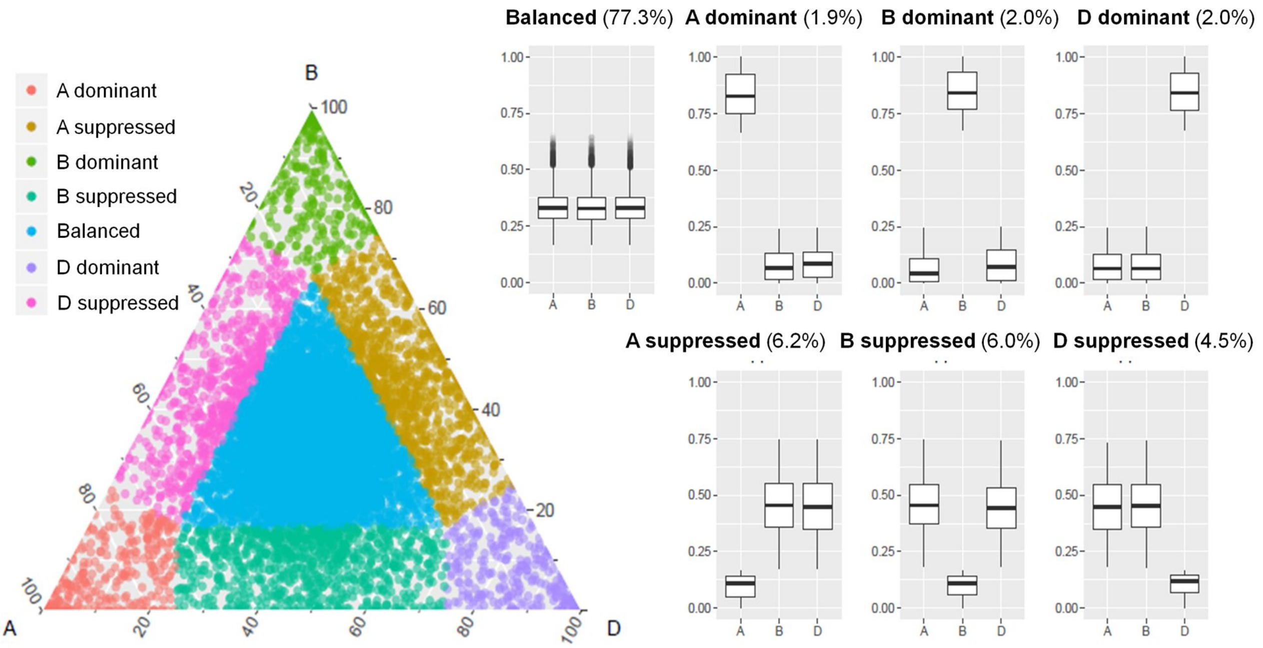

### S2 Fig. Proportion of genes in each homeologs number category

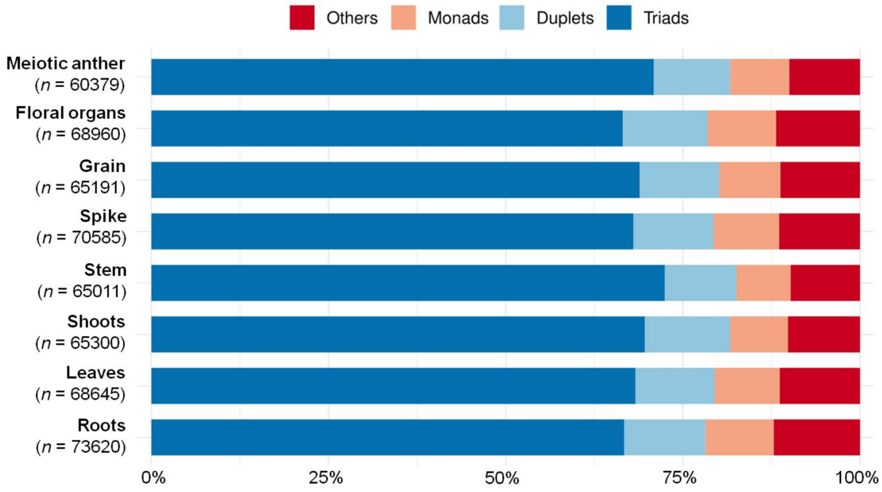

### S4 Fig. Enriched GO terms in the meiosis-related and other tissue-related modules

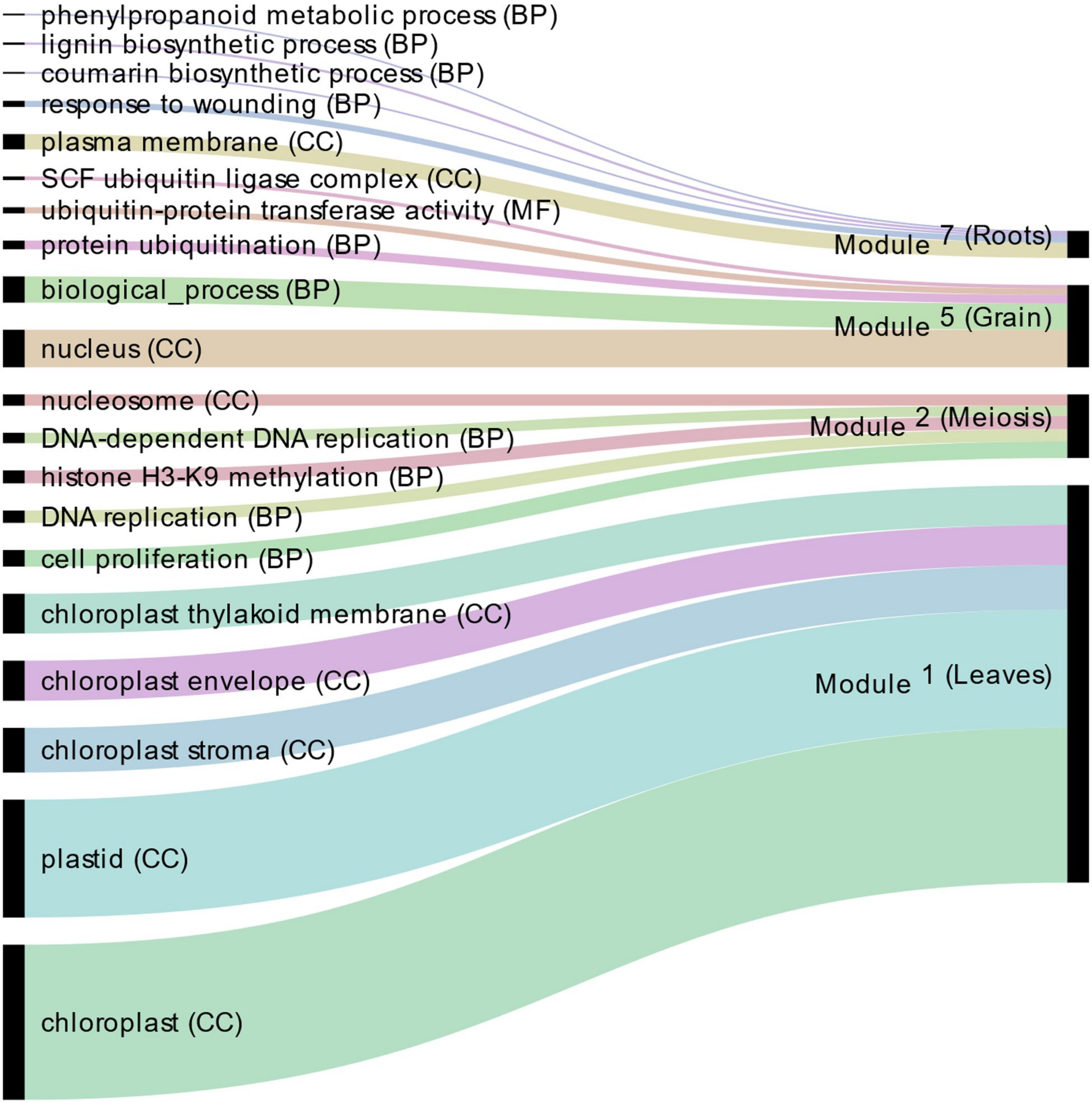
