## Supplementary material for "A co-expression network in hexaploid wheat reveals mostly balanced expression and lack of significant gene loss of homeologous meiotic genes upon polyploidization": S3 Fig. Module-tissue relationship

| | Meiotic anther | Floral organs | Grain | Leaves | Roots | Shoots | Spike | Stem |
|---|---|---|---|---|---|---|---|---|
| ME1 | −0.0014 (1) | −0.12 (1) | −0.59 (1e−11) | 0.72 (5e−20) | −0.29 (0.02) | 0.33 (0.003) | 0.12 (1) | 0.14 (1) |
| ME2 | 0.61 (9e−13) | 0.086 (1) | −0.38 (3e−04) | −0.3 (0.01) | 0.28 (0.03) | −0.13 (1) | −0.07 (1) | 0.011 (1) |
| ME5 | −0.076 (1) | 0.063 (1) | 0.89 (2e−43) | −0.35 (0.001) | −0.42 (2e−05) | −0.19 (0.4) | −0.11 (1) | −0.19 (0.4) |
| ME7 | −0.13 (1) | −0.14 (1) | −0.68 (4e−17) | 0.034 (1) | 0.7 (3e−18) | 0.11 (1) | 0.19 (0.4) | 0.2 (0.3) |
| ME9 | 0.14 (1) | −0.099 (1) | −0.55 (1e−09) | −0.13 (1) | 0.76 (2e−23) | 0.0093 (1) | −0.069 (1) | 0.072 (1) |
| ME11 | 0.68 (1e−16) | 0.38 (3e−04) | 0.06 (1) | −0.42 (3e−05) | −0.43 (2e−05) | −0.23 (0.2) | 0.11 (1) | −0.15 (1) |
| ME13 | 0.016 (1) | −0.19 (0.4) | 0.89 (2e−43) | −0.41 (4e−05) | −0.14 (1) | −0.2 (0.3) | −0.27 (0.03) | −0.16 (0.8) |
| ME17 | −0.53 (7e−09) | −0.19 (0.4) | −0.32 (0.005) | 0.53 (4e−09) | 0.2 (0.3) | 0.25 (0.07) | 0.12 (1) | 0.074 (1) |
| ME19 | 0.46 (1e−06) | 0.081 (1) | 0.63 (1e−13) | −0.43 (2e−05) | −0.43 (2e−05) | −0.23 (0.1) | −0.17 (0.6) | −0.22 (0.2) |
| ME20 | −0.25 (0.07) | −0.19 (0.4) | −0.58 (3e−11) | 0.45 (3e−06) | 0.57 (1e−10) | 0.18 (0.5) | −0.07 (1) | 0.033 (1) |
| ME21 | −0.37 (5e−04) | −0.32 (0.005) | 0.22 (0.2) | 0.65 (1e−14) | −0.48 (5e−07) | 0.27 (0.03) | −0.069 (1) | 0.021 (1) |
| ME23 | −0.29 (0.02) | −0.045 (1) | 0.79 (7e−27) | −0.17 (0.7) | −0.21 (0.3) | −0.061 (1) | −0.24 (0.1) | −0.14 (1) |
| ME25 | 0.65 (9e−15) | 0.41 (4e−05) | −0.48 (4e−07) | −0.2 (0.4) | −0.29 (0.02) | −0.14 (1) | 0.31 (0.008) | 0.016 (1) |
| ME26 | 0.17 (0.6) | −0.0083 (1) | 0.68 (9e−17) | −0.49 (2e−07) | −0.23 (0.1) | −0.21 (0.3) | −0.12 (1) | −0.084 (1) |
| ME27 | −0.25 (0.08) | −0.23 (0.1) | 0.74 (8e−22) | −0.46 (2e−06) | 0.22 (0.2) | −0.21 (0.3) | −0.12 (1) | −0.092 (1) |
| ME28 | 0.52 (2e−08) | 0.15 (1) | 0.24 (0.1) | −0.57 (8e−11) | −0.037 (1) | −0.26 (0.06) | −0.088 (1) | −0.077 (1) |
| ME29 | −0.31 (0.009) | 0.17 (0.7) | 0.69 (3e−17) | −0.25 (0.07) | −0.33 (0.003) | −0.14 (1) | 0.12 (1) | −0.18 (0.5) |
| ME32 | −0.48 (3e−07) | −0.35 (0.001) | 0.85 (1e−34) | −0.12 (1) | −0.026 (1) | −0.032 (1) | −0.15 (0.9) | −0.11 (1) |
| ME33 | 0.19 (0.4) | −0.15 (1) | 0.62 (2e−13) | −0.55 (1e−09) | 0.064 (1) | −0.25 (0.07) | −0.19 (0.4) | −0.08 (1) |
| ME36 | −0.18 (0.6) | −0.18 (0.6) | 0.83 (7e−32) | −0.23 (0.2) | −0.21 (0.3) | −0.15 (0.9) | −0.17 (0.6) | −0.13 (1) |
| ME38 | 0.053 (1) | 0.01 (1) | 0.64 (2e−14) | −0.21 (0.2) | −0.23 (0.1) | −0.033 (1) | −0.37 (5e−04) | −0.15 (1) |
| ME39 | 0.034 (1) | −0.34 (0.002) | 0.5 (6e−08) | −0.24 (0.09) | 0.13 (1) | −0.14 (1) | −0.27 (0.03) | 0.018 (1) |
| ME41 | 0.73 (3e−20) | 0.19 (0.4) | 0.057 (1) | −0.37 (4e−04) | −0.28 (0.02) | −0.22 (0.2) | −0.031 (1) | −0.12 (1) |
| ME45 | −0.34 (0.002) | −0.24 (0.1) | −0.073 (1) | 0.68 (5e−17) | −0.32 (0.005) | 0.29 (0.02) | 0.035 (1) | 0.021 (1) |
| ME47 | −0.31 (0.008) | −0.19 (0.4) | 0.8 (4e−28) | 0.093 (1) | −0.47 (6e−07) | 0.018 (1) | −0.14 (1) | −0.15 (1) |
| ME48 | −0.53 (6e−09) | 0.054 (1) | 0.7 (3e−18) | −0.1 (1) | −0.18 (0.5) | −0.083 (1) | −0.00039 (1) | −0.13 (1) |
| ME50 | −0.028 (1) | −0.47 (9e−07) | 0.78 (2e−25) | −0.18 (0.5) | −0.071 (1) | −0.049 (1) | −0.34 (0.002) | −0.076 (1) |
| ME55 | −0.65 (7e−15) | −0.28 (0.03) | 0.56 (5e−10) | −0.28 (0.03) | 0.33 (0.004) | −0.098 (1) | 0.13 (1) | 0.044 (1) |
| ME57 | −0.55 (6e−10) | −0.062 (1) | 0.58 (6e−11) | 0.069 (1) | −0.086 (1) | 0.016 (1) | −0.081 (1) | −0.14 (1) |
| ME60 | −0.48 (5e−07) | −0.2 (0.3) | −0.1 (1) | 0.71 (3e−19) | −0.19 (0.4) | 0.36 (9e−04) | −0.065 (1) | 0.036 (1) |
| ME61 | −0.18 (0.5) | −0.18 (0.5) | 0.62 (3e−13) | 0.2 (0.3) | −0.62 (4e−13) | 0.062 (1) | −0.073 (1) | −0.07 (1) |
| ME64 | 0.23 (0.2) | −0.066 (1) | −0.49 (2e−07) | −0.21 (0.2) | 0.85 (3e−34) | −0.071 (1) | −0.13 (1) | −0.055 (1) |

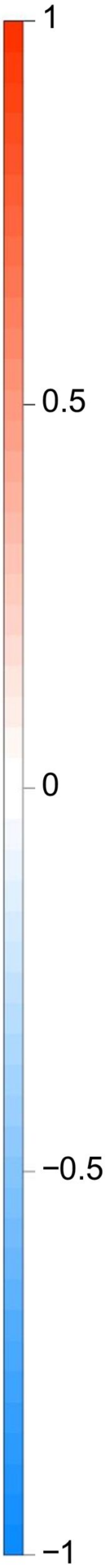
